# Manipulation of the *brown glume and internode 1* gene leads to alterations in lignified tissue coloration, lignification, and pathogen resistance in wheat

**DOI:** 10.1101/2024.10.11.617803

**Authors:** Lei Hua, Rui Song, Xiaohua Hao, Jing Zhang, Yanna Liu, Jing Luo, Xiaopeng Ren, Hongna Li, Guiping Wang, Shams ur Rehman, Jiajie Wu, Daolin Fu, Yuxiu Dong, Xiaodong Wang, Chaozhong Zhang, Shisheng Chen

## Abstract

Lignin is a crucial component of the cell wall, providing mechanical support and protection against biotic and abiotic stresses. However, little is known about wheat lignin-related mutants and their roles in pathogen defense. In this study, we identified an ethyl methanesulfonate (EMS)-derived *Aegilops tauschii* mutant named *brown glume and internode 1* (*bgi1*), which exhibits reddish-brown pigmentation in various tissues, including internodes, spikes, and glumes. Using map-based cloning and single nucleotide polymorphism (SNP) analysis, we identified *AET6Gv20438400* (*BGI1*) as the leading candidate gene, encoding the TaCAD1 protein. The mutation occurred in the splice acceptor site of the first intron, resulting in a premature stop codon in *BGI1*. We validated the function of *BGI1* using loss-of-function EMS and gene editing knockout mutants, both of which displayed reddish-brown pigmentation in lignified tissues. *BGI1* knockout mutants exhibited reduced lignin content and shearing force relative to wild type, while *BGI1* overexpression transgenic plants showed increased lignin content and enhanced disease resistance against common root rot and *Fusarium* crown rot. We confirmed that BGI1 exhibits CAD activity both *in vitro* and *in vivo*, playing an important role in lignin biosynthesis. *BGI1* was highly expressed in the stem and spike and localized in the cytoplasm. Transcriptome analysis revealed the regulatory networks associated with *BGI1*. Finally, we demonstrated that BGI1 interacts with TaPYL-1D, potentially involved in the abscisic acid signaling pathway. The identification and functional characterization of *BGI1* significantly advance our understanding of CAD proteins in lignin biosynthesis and plant defense against pathogen infection in wheat.

## Introduction

Bread wheat (*Triticum aestivum* L.) is a crucial staple crop worldwide, contributing about 20% of the world’s caloric and protein intake for the human population. However, wheat production is threatened by various factors, including diseases and lodging. Common root rot (CRR) caused by *Bipolaris sorokiniana* and *Fusarium* crown rot (FCR) caused by multiple *Fusarium* species are prevalent diseases in many arid and semi-arid cropping regions worldwide, such as Australia, the United States, Canada, China, and South Africa (Bozoğlu et al., 2022; Kazan and Gardiner, 2018). These two diseases are major concerns in regions with extensive wheat-maize rotation and straw returning practices, as seen in China (Su et al., 2021). In addition to diseases, lodging is also a significant limiting factor for wheat production, reducing grain yield and causing several knock-on effects, including decreased grain quality and increased drying costs (Berry and Spink, 2012). Enhancing lignin content to improve lodging resistance is a crucial breeding objective (Zhang et al., 2016).

Lignin is a complicated and heterogeneous aromatic polymer that forms the secondary cell wall together with cellulose and hemicellulose. Lignin plays an important role in providing mechanical support, promoting water conductance, and offering protection against biotic and abiotic stresses during plant growth and development (Barros et al., 2015; Gallego-Giraldo et al., 2020). Lignin is gaining increasing attention due to its crucial role in enhancing both abiotic stress tolerance and pathogen resistance (Lee et al., 2019; Rong et al., 2016; Xu et al., 2020). The synthesis of lignin involves the polymerization of monolignols, which are produced through the phenylpropanoid pathway and subsequently undergo hydroxylation and methylation processes (Boerjan et al., 2003; Ralph et al., 2004). Polymerization of the three main monolignols (*p*-coumaryl, coniferyl, and sinapyl alcohols) resulted in the formation of various monomers, namely *p*-hydroxyphenyl (H), guaiacyl (G), and syringyl (S), respectively (Vanholme et al., 2010).

Cinnamyl alcohol dehydrogenase, also known as CAD, is an enzyme that plays a crucial role in the lignin biosynthesis pathway. It is responsible for catalyzing the nicotinamide adenine dinucleotide phosphate (NADPH)-dependent reduction of cinnamaldehydes to cinnamyl alcohol, the final step in monolignol biosynthesis before polymerization in the cell wall (Goffner et al., 1992; Vanholme et al., 2010). Disruption of CAD activity significantly increases the incorporation of cinnamaldehyde into the lignin polymer (Liu et al., 2021c). Plants with impaired CAD function may exhibit delayed flowering, reduced plant height, and a characteristic brownish-red to tan pigmentation in the leaf midrib, particularly in C4 grasses (Chen et al., 2012; Liu et al., 2021c; Trabucco et al., 2013). Previous studies indicated that a high amount of coniferyl aldehyde is the chromophoric origin of the reddish-brown coloration (Tsai et al., 1998). The complete genome sequencing and annotation have enabled researchers to determine the number of *CAD* genes in various species, which has led to their classification into three classes (Barakat et al., 2009). Class I CADs exhibit a highly conserved primary sequence structure across most species and are involved in lignin deposition in the secondary cell walls during plant growth and development (Park et al., 2018). Class II CADs represent a more extensive and diverse group of CAD isoforms, known to be associated with stress resistance (Rong et al., 2016). Class III CAD members may show redundancy with Class I CADs, yet their precise functions remain unclear (Peracchi et al., 2024; Xu et al., 2011).

In *Arabidopsis*, although nine *CAD* genes have been identified in the published reference genome (Sibout et al., 2005), only three (*AtCAD1*, *AtCAD4*, and *AtCAD5*) have been validated as key enzymes involved in monolignol biosynthesis (Rong et al., 2016). Similarly, among the 12 *CAD* genes identified in the rice genome, *OsCAD2* has been directly linked to lignin biosynthesis (Zhang et al., 2006), while *OsCAD7* plays a role in culm mechanical strength (Li et al., 2009). Recently, in silico analysis of the hexaploid wheat genome have revealed 47 high-confidence *TaCAD* gene copies (Peracchi et al., 2024). Among these *TaCAD* genes, *TaCAD1* was speculated to be involved in lodging resistance (Chen et al., 2021; Ma, 2010), and *TaCAD12* was found to contribute to host resistance against sharp eyespot (Rong et al., 2016). However, the functionality of neither gene has been experimentally verified in wheat. The challenges posed by the size and complexity of the bread wheat genome have historically impeded the identification and characterization of *TaCAD* genes. Currently, there is a lack of data on lignin-related *CAD* mutants or overexpression in wheat. The present study addresses this gap by using a lignin-deficient mutant called *brown glume and internode 1* (*bgi1*) identified from an EMS-mutagenized library of the diploid *Aegilops tauschii* accession PI 511383. The *bgi1* mutant plants exhibit a reddish-brown pigmentation in various tissues, including internodes, spikes, and glumes. Through map-based cloning and single nucleotide polymorphism (SNP) analysis, we isolate the *BGI1* gene, which encodes the TaCAD1 protein. We validate the function of the cloned candidate gene using loss-of-function EMS and gene editing knockout mutants. Furthermore, we investigate the regulatory mechanisms determining the role of BGI1 in lignin biosynthesis and demonstrate that *BGI1* overexpression increases lignin content and enhances resistance against both CRR and FCR.

## Results

### Characterization of the *bgi1* mutant

We identified an M_2_ mutant line, M326 (*bgi1*), derived from an EMS-mutagenized population of the *Ae. tauschii* accession PI 511383. A morphological comparison between the *bgi1* mutant and wild type (WT) plants revealed that *bgi1* mutants exhibited normal development under greenhouse conditions (Fig. 1a). However, at the heading stage, various parts of the *bgi1* mutant plant, including the internode, spike, spike rachilla, glume, and lemma, exhibited distinctive reddish-brown pigmentation (Figs. 1b-g). The reddish-brown pigmentation became evident at the jointing stage and reached its peak at the heading stage. The spikes showed a reddish-brown coloration (Fig. 1c), and the node of the spike rachilla facing the spikelet displayed the most vivid reddish-brown hue (Fig. 1d). Interestingly, the interior of the glume displayed a deeper hue compared to the exterior (Figs. 1e, f).

**Figure 1.**
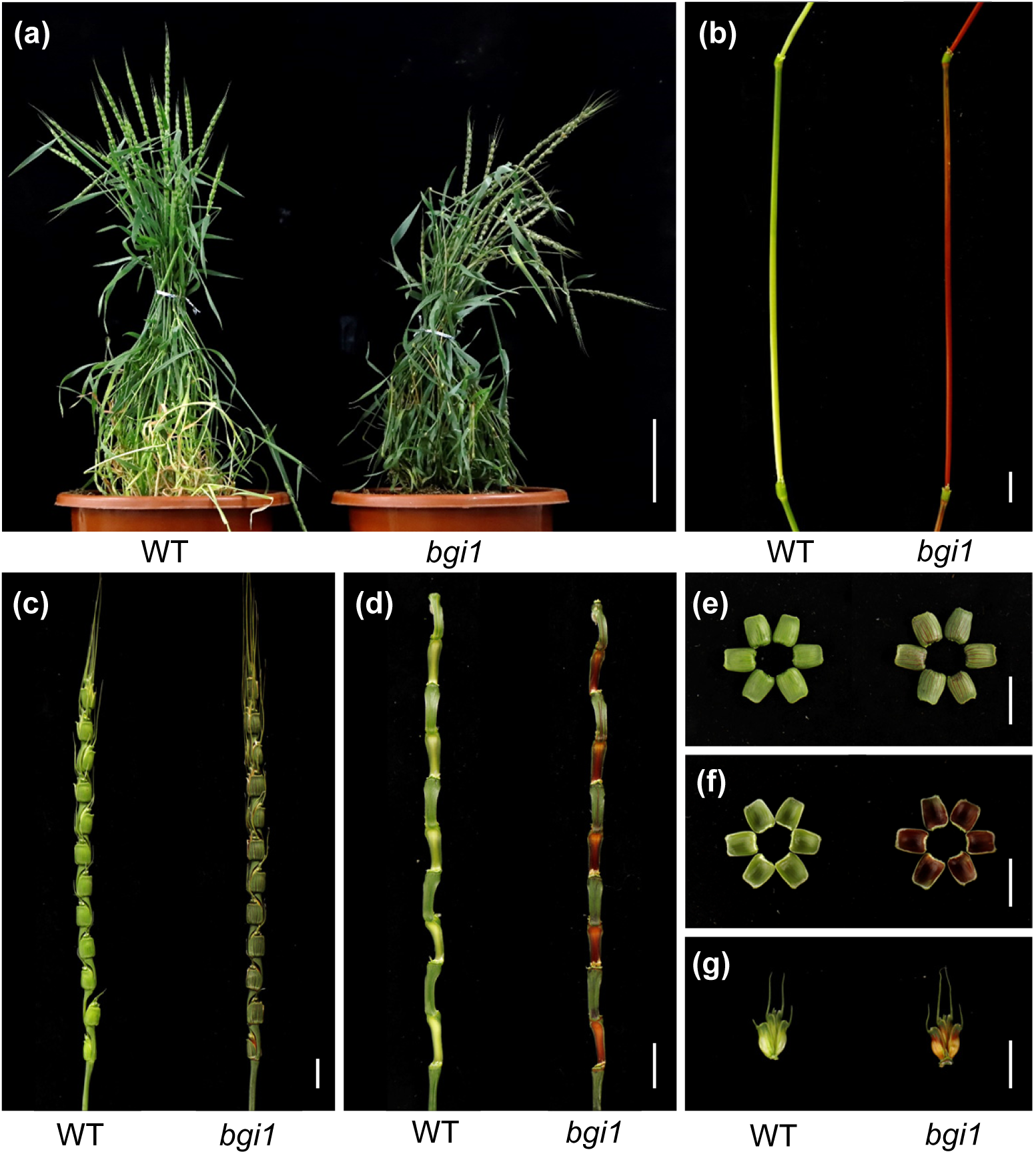
Phenotypic comparison between the wild type (WT) and the *bgi1* mutant. (a-g) Close-up views of the plant (a), internode (b), spike (c), spike rachilla (d), outer of the glume (e), inner of the glume (f), and the spikelet after removing the glumes (g) between WT and the *bgi1* mutant at the heading stage. Scale bars, 10 cm in (a) and 1 cm in (b-g).

Crossing the *bgi1* mutant with the WT (PI 511383) produced F_1_ plants that uniformly exhibited the WT phenotype. Within a subset of 94 F_2_ plants derived from this cross, 68 plants displayed normal coloration, while 26 plants exhibited reddish-brown glumes and internodes, which fits the expected 3:1 segregation ratio for a single recessive gene (χ*^2^* = 0.35, *P* = 0.55).

### Map-based cloning of *bgi1*

To map the causal gene, two F_2_ mapping populations were generated: one comprising 182 F_2_ plants derived from the M326 × AL8/78 cross, and the other comprising 1,872 F_2_ plants from the M326 × PI 511383 cross. Through RNA sequencing and SNP calling, 225 high-confidence EMS-induced SNPs were identified between M326 and PI 511383. Bulked segregant RNA sequencing (BSR-Seq) analysis revealed 18 SNPs on chromosome 6D, ranging from 40.2 to 324.1 Mb (AL8/78 Aet v4.0; Table S1), which are significantly associated with the phenotype in the M326 × PI 511383 population. This finding indicates that the *bgi1* gene is located on chromosome 6D.

To confirm the mapping location, seven SNPs within the candidate region on chromosome 6D (Table S1) were selected for developing D-genome specific cleaved amplified polymorphic sequence (CAPS) or kompetitive allele-specific PCR (KASP) markers (Table S2). Genotyping of 94 F_2_ plants from the M326 × PI 511383 cross with these markers pinpointed the *bgi1* gene within a 2.2 cM interval flanked by CAPS markers *pkw1225* and *pkw2754* (Fig. 2b). In the mapping population derived from the M326 × AL8/78 cross, ten PCR markers were developed (Table S2), mapping *bgi1* within a 1.65 cM interval flanked by markers *ucw139* and *ucw239* (Fig. S1). Notably, both populations exhibited recombination suppression in regions approximately from 157 to 239 Mb of chromosome 6D.

**Figure 2.**
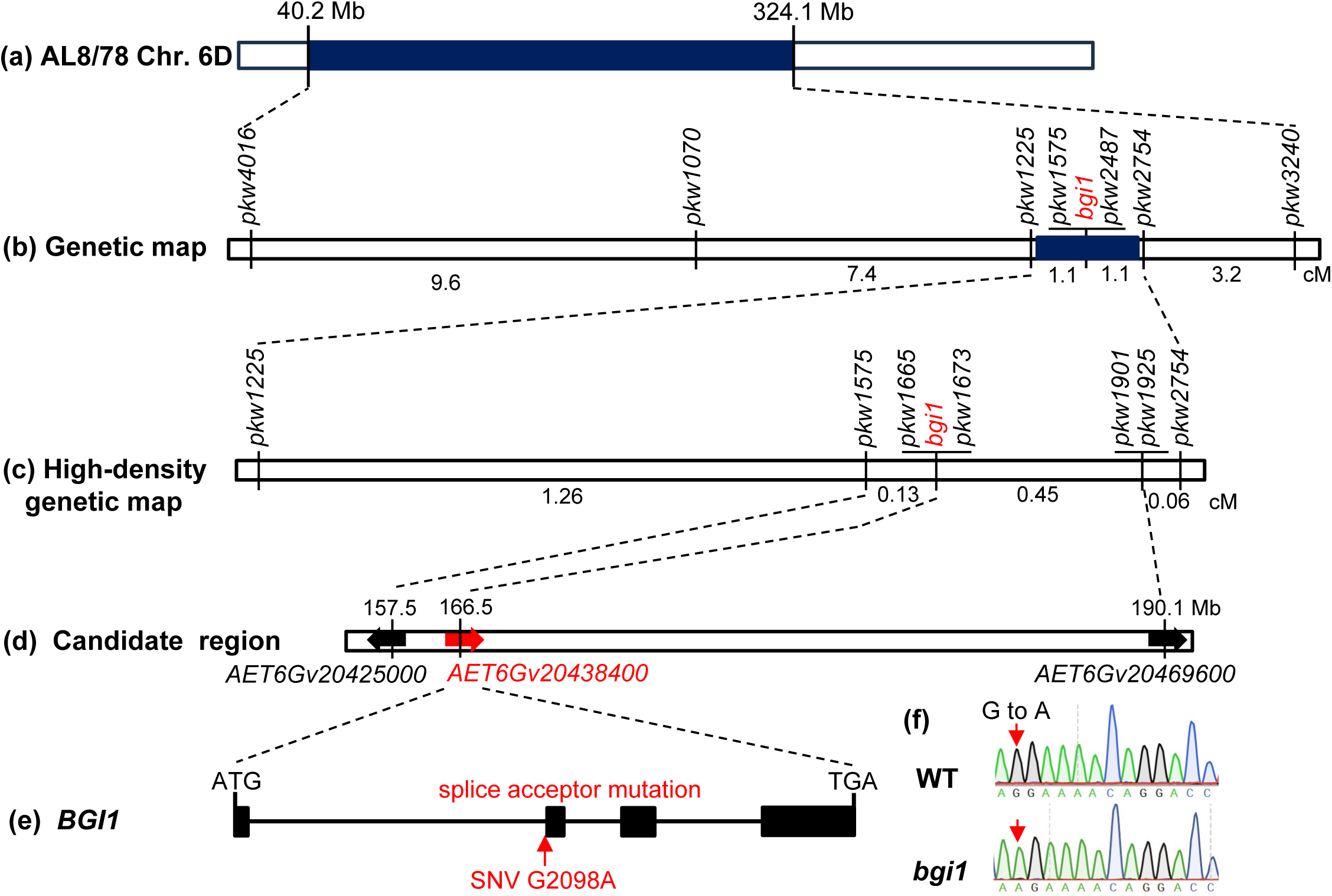
Map-based cloning of the causal gene *bgi1*. (a) Genetic mapping of *bgi1* on chromosome 6D by BSR-seq. Genomic region containing *bgi1* was highlighted in blue. (b) Genetic map based on 94 F_2_ plants from the M326 × PI 511383 cross and seven molecular markers. (c) High-density genetic map based on 1,872 F_2_ plants and another seven molecular markers. (d) Physical map of *bgi1* in the sequenced reference genome of *Ae. tauschii* AL8/78 (Aet v4.0). Among the 213 annotated high-confidence genes within the mapping region (Table S3), only the candidate gene is highlighted and marked with a red arrow. (e) Gene structure of the candidate gene *AET6Gv20438400* (*BGI1*). Black rectangles and lines represent exons and introns, respectively; the mutated site was marked in red. (f) Sequencing chromatograms showing the G to A polymorphism between two parents. The mutated nucleotide is indicated by a red arrow. cM, centimorgans; Mb, megabases.

To further narrow down the mapping region of *bgi1*, an additional 1,778 F_2_ plants from the M326 × PI 511383 cross were screened for recombinants between markers *pkw1225* and *pkw2754*. Using these recombinants and four newly developed markers (Table S2), the *bgi1* gene was further mapped to a 0.58 cM region flanked by markers *pkw1575* and *pkw1901*, and was completely linked to markers *pkw1665* and *pkw1673* (Fig. 2c). The recombination rate was significantly reduced within the *bgi1* candidate region, likely due to its proximity to the centromere.

The 0.58 cM candidate region encompasses a 32.6-Mb region (157.5-190.1 Mb) in the *Ae. tauschii* reference genome (AL8/78 Aet v4.0), containing 213 annotated high-confidence genes (Table S3). RNA sequencing results identified only two high-confidence SNPs (G166511490A and G167343806A) between M326 and PI 511383 within the candidate region (Table S1). The SNP, G167343806A, is located within the 5’ untranslated region (UTR) or the intron of *AET6Gv20439300* (depending on the alternative splice forms), which encodes a glycerophosphodiester phosphodiesterase protein. Analysis of published RNA-seq data from the wheat expVIP database (https://www.wheat-expression.com/) revealed very low expression of this gene in the stem and spike (Fig. S2a), suggesting it is unlikely to be the *bgi1* gene.

The other SNP, G166511490A, is located within the candidate gene *AET6Gv20438400*, where it acts as a splice acceptor mutation (Fig. 2e). Expression profile analysis revealed that *AET6Gv20438400* is highly expressed in various wheat tissues, particularly in the stem and spike (Fig. S2b). Sanger sequencing confirmed the presence of a mutation (G > A) at the splicing acceptor site of the first intron of the candidate gene *AET6Gv20438400*, leading to premature termination of protein (Fig. 2f).

*AET6Gv20438400* (*BGI1*) spans 4,233 bp from the start to the stop codon, comprising four exons and three introns (Fig. 2e). The complete coding sequence of 1,083 bp encodes a predicted protein of 360 amino acids. *BGI1* is orthologous to the maize gene *BM1* (Halpin et al., 1998), the rice gene *GH2* (Zhang et al., 2006), the sorghum gene *BMR6* (Li et al., 2015), and the *Brachypodium distachyon* gene *BdCAD1* (Bouvier d’Yvoire et al., 2013). Phylogenetic analysis showed that BGI1 was grouped in a clade with these known CAD proteins (Fig. S3). Mutations in these *CAD* genes have been associated with pigmentation changes in lignified tissues (Bouvier d’Yvoire et al., 2013; Halpin et al., 1998; Li et al., 2015; Zhang et al., 2006). These results make *BGI1* a highly promising candidate.

### Loss-of-function EMS mutations in *BGI1* homeologs showed reddish-brown phenotypes

Within the tetraploid and hexaploid wheat genomes, the homeologs of BGI1 exhibit a similarity exceeding 98.9% (Fig. S4), suggesting potential functional redundancy. To confirm the casual relationship between *BGI1* and observed phenotypic alterations, two mutant lines, K3288 and K3088, were selected from a sequenced mutant population of the tetraploid wheat variety Kronos (Krasileva et al., 2017). The mutant line K3288 harbors a single nucleotide mutation in the fourth exon of the A-genome copy (C3623T), resulting in a premature stop codon (Q312*; Fig. 3a). Similarly, the mutant line K3088 carries a G42A mutation in the first exon the B-genome copy, leading to a premature stop codon (W14*; Fig. 3a). Using genome-specific markers *K3288-A1* and *K3088-B1* (Table S2), we identified mutant plants homozygous for either the *bgi-A1* or *bgi-B1* allele (Fig. 3b). In greenhouse experiments, neither *bgi-A1* nor *bgi-B1* homozygous plants exhibited reddish-brown phenotypes. Subsequently, we crossed the K3288 and K3088 mutants, and selected F_2_ plants homozygous for all four possible combinations of the *BGI1* homeologs-WT, *bgi-A1*, *bgi-B1*, and *bgi1* double mutants for further evaluation. In this experiment, only the *bgi1* double mutant plants displayed reddish-brown pigmentation at the heading stage, while neither the *bgi-A1* nor the *bgi-B1* mutants exhibited phenotypic alterations (Figs. 3c-i). These results demonstrate that *BGI-A1* and *BGI-B1* genes exhibit functional redundancy in tetraploid wheat.

**Figure 3.**
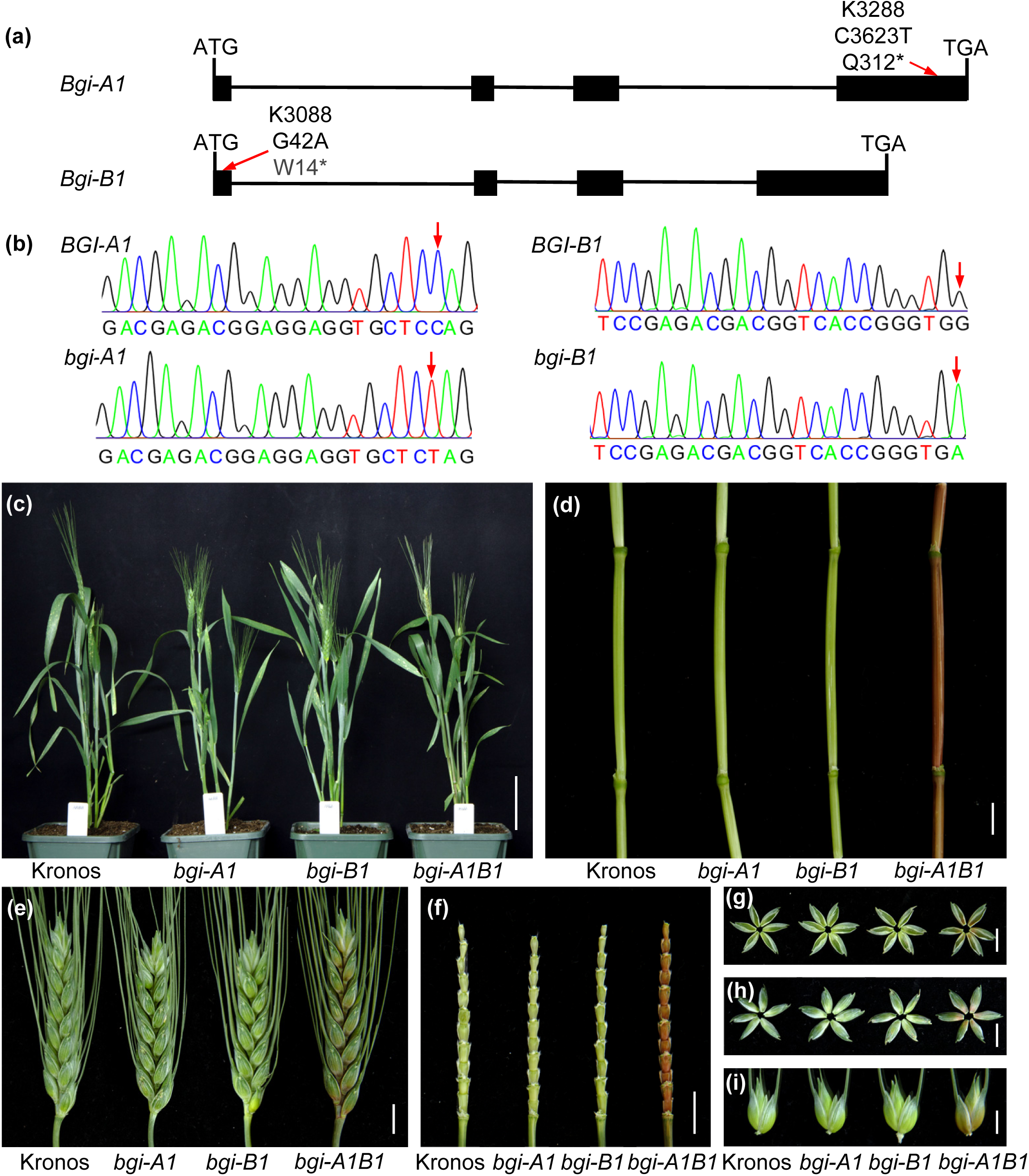
Functional validation of *BGI1* using EMS mutants. (a) Gene structure of the *BGI1* homeologs in tetraploid wheat variety Kronos. Exons are represented by black boxes and introns by black lines. EMS mutation sites are indicated by red arrows. (b) Sequencing chromatograms showing nucleotide mutations in *BGI-A1* and *BGI-B1*. The mutant nucleotides are highlighted by red arrows. (c-i) Phenotypic comparison of the plant (c), internode (d), spike (e), spike rachilla (f), outer glume (g), inner glume (h), and spikelet with glumes removed (i) among Kronos (WT), single-gene mutants (*bgi-A1* and *bgi-B1*), and double mutants (*bgi-A1B1*) at the heading stage. Scale bars, 10 cm in (c) and 1 cm in (d-i).

In the sequenced EMS-mutagenized population of hexaploid wheat variety Cadenza (Krasileva et al., 2017), we selected truncation mutations for the A, B, and D genome homeologs of *BGI1* (Fig. S5a). To generate loss-of-function EMS mutants in hexaploid wheat background, we combined the splice site mutations in the A-and B-genome homeologs (*bgi-A1* and *bgi-B1*) with a premature stop codon mutation (Q322*) in the D-genome homeolog (*bgi-D1*) by crossing and selection in the F_2_ generation. The procedure for generating single-gene mutants, double mutants, and the *bgi1* triple mutant is detailed in Figure S6. Phenotypic analysis revealed that only the *bgi1* triple mutant plants exhibited a reddish-brown phenotype at the heading stage, and none of the single or double mutants showed phenotypic alterations (Figs. S5b-h). These results confirm the redundant roles of these three homeologs in determining the coloration of glumes and internodes.

### Validation of *BGI1* using barley stripe mosaic virus (BSMV)-sgRNA-based gene editing

To validate the phenotypic observations associated with the EMS-induced *bgi1* mutant, we employed a BSMV-sgRNA-based gene editing approach (Li et al., 2021) to generate independent edited wheat plants. We designed a sgRNA that specifically targeting a conserved region in the first exon, enabling the simultaneous knockout of all three *BGI1* homeologs (Fig. 4a). The wheat transgenic line expressing the *Cas9* gene in the Bobwhite genetic background (*Cas9*-transgenic Bobwhite) was infected with BSMV vectors. Ten out of eleven infected M_0_ plants in *Cas9*-transgenic Bobwhite were edited, as confirmed by DNAs extraction from flag leaves and genotyping with the CAPS marker *HL553* (digested with *Psh*AI; Figs. S7a, b).

**Figure 4.**
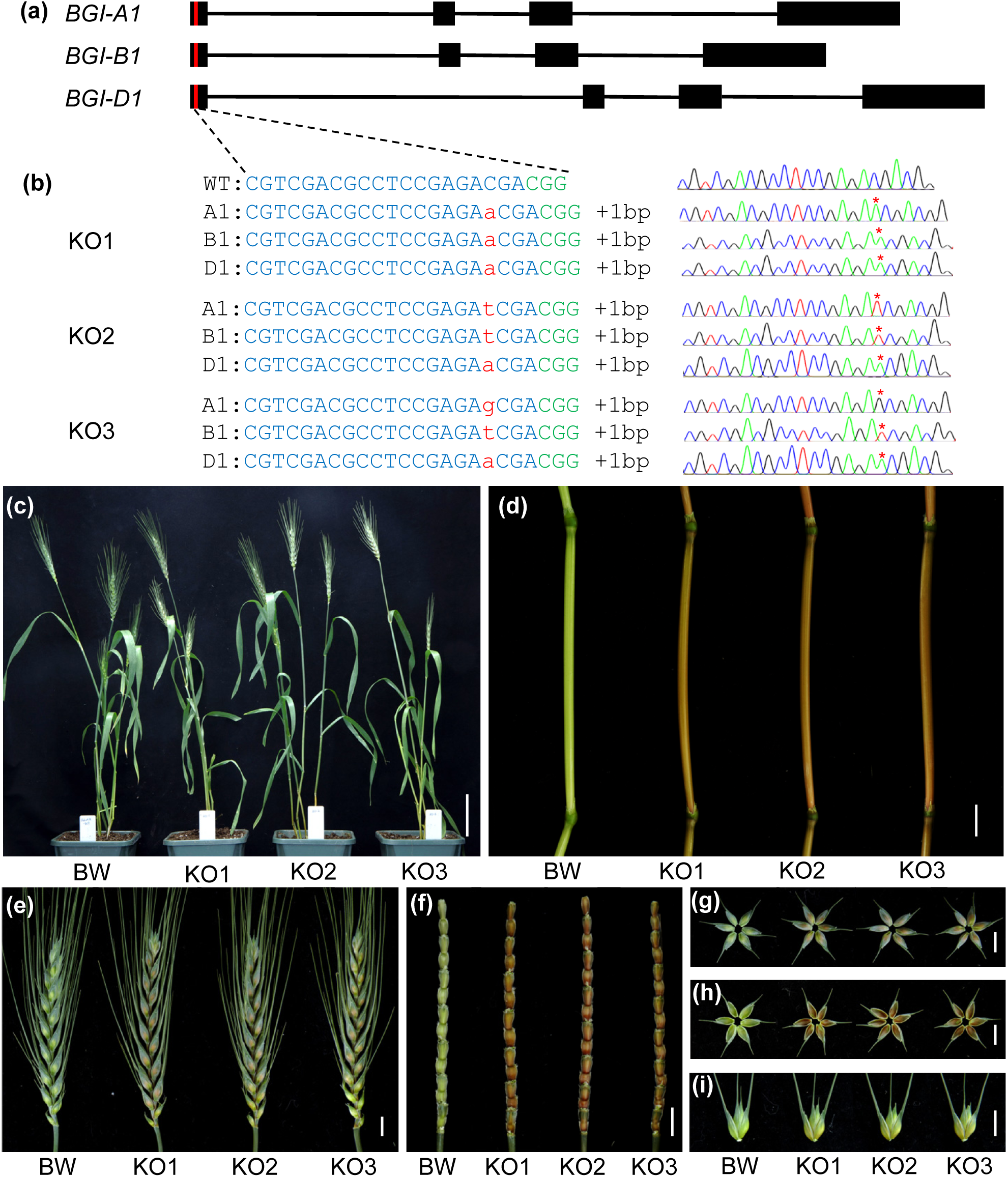
Validation of *BGI1* using BSMV-sgRNA-based gene editing. (a) Gene structure of the *BGI1* homeologs in Bobwhite. Exons are represented by black boxes and introns by black lines. Red boxes indicate the target sites. (b) Sequencing chromatograms showing the induced polymorphisms between WT (Bobwhite) and selected BSMV-induced editing mutant lines (KO1-KO3). The mutated nucleotides are highlighted in red and the CGG PAM sequences are marked in green. *, mutated nucleotides. (c-i) Phenotypic comparison of the plant (c), internode (d), spike (e), spike rachilla (f), outer glume (g), inner glume (h), and spikelet with glumes removed (i) between Bobwhite (BW) and KO mutant lines at the heading stage. Scale bars, 10 cm in (c) and 1 cm in (d-i).

Subsequently, we genotyped 99 M_1_ plants derived from the selected M_0_ edited plants using the genome-specific markers *HL554*, *HL555*, and *HL556* (Table S2). Sanger sequencing revealed mutations in 87 of the 99 M_1_ plants tested, yielding a mutation frequency of 87.9% (Table S4). Among these 87 edited M_1_ plants, 41 harbored mutations in all three *BGI1* homeologs, with three being homozygous knockout (KO) mutants wherein six alleles were edited concurrently (referred to as KO1, KO2, and KO3; Table S4).

These three independent homozygous KO mutant lines carry insertions of an “A” or “T” or “G” at position 25 from the ATG (3 bp upstream of the CGG PAM site) in all three *BGI1* homeologs (Fig. 4b). These frameshift insertions alter more than 97.2% of the protein sequences, resulting in loss-of-function in all three homeologs. Phenotypic investigations showed that single and double mutant plants exhibit no phenotypic changes (Fig. S8). In contrast, the homozygous KO mutant lines KO1, KO2, and KO3 exhibited reddish-brown pigmentation in the internode, spike, spike rachilla, glume, and lemma at the heading stage (Figs. 4c-i).

Taken together, the genetic mapping, SNP analysis, the loss-of-function EMS mutations, and BSMV-sgRNA-based gene editing evidently demonstrated that *BGI1* is the causal gene responsible for the *bgi1* mutant phenotype.

### Disrupting or overexpressing *BGI1* results in significant changes in lignin content and composition

To investigate the cellular basis of the reddish-brown coloration observed in the mutant line KO1, we performed histological examinations of transverse sections from internodes at the heading stage. Reddish-brown pigmentation was predominantly localized to lignified tissues, mainly in the mechanical tissue (MT) and vascular bundle (VB) regions (Figs. 5a, d). Subsequent staining with Wiesner reagent revealed pronounced variations between WT and KO1 mutant samples (Figs. 5b, e). The cross-sections of WT displayed a purple hue from the epidermal layer to the MT and VB regions (Fig. 5b), whereas the KO1 mutant exhibited significantly reduced staining (Fig. 5e), indicating a lower level of lignin accumulation in the mutant internodes. Mäule staining (Meyer et al., 1998), which differentiates G residues as yellow and S residues as red, was used to assess lignin composition. The strong red coloration of lignified tissues observed in WT was significantly reduced in the KO1 mutant samples (Figs. 5c, f), suggesting that KO1 impedes the formation of S residues in lignified stem tissues. Apart from the differences mentioned, no other significant differences were observed between WT and KO1 plants across the seven agronomic traits examined under greenhouse conditions (Fig. S9).

**Figure 5.**
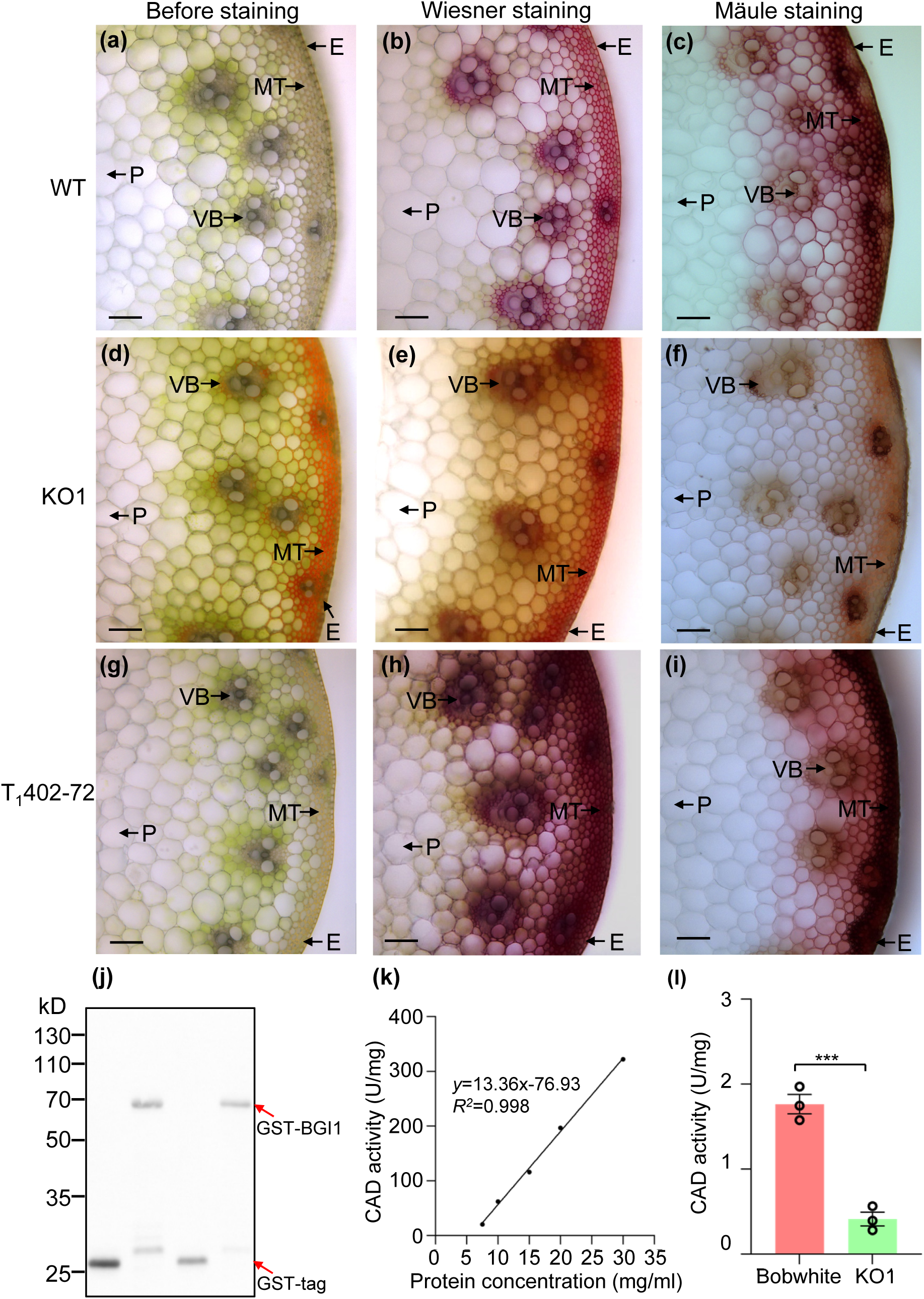
B*G*I1 is involved in lignin biosynthesis. (a-i) Cross-sections before staining, after Wiesner staining, and after Mäule staining from WT (Bobwhite; a-c), KO1 mutant plants (d-f), and T_1_402-72 transgenic plants (g-i). Hand-cut cross-sections of second internodes at the heading stage were used. E, epidermis; MT, mechanical tissue; VB, vascular bundle; P, parenchyma. Scale bars, 100 μm. (j) Western blot analysis of glutathione S-transferase (GST)-BGI1 protein using anti-GST antibodies. The molecular weights of purified GST and GST-BGI1 proteins are 26 kD and 66 kD, respectively, as indicated by red arrows. (k) Correlation between BGI1’s enzymatic activity and the concentration of the recombinant GST-BGI1 protein. The substrates, coniferyl aldehyde and NADPH, were used. The reaction mixtures were incubated in a water bath at 25℃ for 5 minutes, after which CAD activity was measured at OD_340_. (l) Comparison of CAD activities between WT and the KO1 mutant plants. Error bars are standard errors of the means (n = 3). ***, *P* < 0.001.

To determine the impact of *BGI1* overexpression on lignin content, we generated nine independent T_0_ transgenic plants overexpressing *BGI1* driven by the maize ubiquitin (UBI) promoter. Transcript levels of *BGI1* were significantly higher (*P* < 0.001) in all transgenic plants compared to the Fielder control (Fig. S10). When stained with Wiesner reagent, transverse sections of plants from transgenic family T_1_402-72 displayed significantly increased staining relative to WT (Fig. 5h), indicating higher lignin content. When subjected to Mäule staining, the MT and VB regions exhibited intense red staining in the T_1_402-72 plants (Fig. 5i), suggesting that *BGI1* overexpression enhances the formation of S units in the stems.

To quantitatively evaluate lignin content, Klason lignin analyses were carried out on dried internodes to determine the lignin content in WT, KO1, and OE transgenic plants. The total lignin content and Klason lignin levels were significantly reduced in KO1 compared to WT, while both were significantly increased in T_1_402-72 and T_1_402-73 transgenic plants (*P* < 0.05; Fig. S11). At the heading stage, the penultimate internodes from WT, KO1, and OE transgenic plants were cut into equal-length segments (∼10 cm) for shearing force measurement. KO1 mutant plants exhibited a significant reduction in shearing force compared to WT plants, whereas OE transgenic plants showed an increased shearing force relative to WT (Fig. S12).

### Overexpression of *BGI1* enhances resistance to both CRR and FCR

Lignin accumulation in cell walls can establish mechanical barriers to pathogen invasion and has been associated with plant defense against pathogen infection (Rong et al., 2016; Tronchet et al., 2010). Based on the observed high accumulation of lignin in the stems of *BGI1*-OE transgenic lines, we hypothesized that the plant resistance to stem base rot diseases such as CRR and FCR will be enhanced by overexpression of *BGI1*. To test this hypothesis, we challenged independent OE transgenic lines and the Fielder control with CRR and FCR in growth chambers. All T_1_ transgenic plants containing the transgene exhibited increased resistance against both CRR and FCR compared to Fielder (Figs. 6a, b). The average lesion area in OE transgenic lines was significantly smaller (*P* < 0.001) than in the Fielder control (Figs. 6c, d). A negative correlation was observed between *BGI1* overexpression levels and average lesion area (*R* = -0.76). These findings demonstrate that *BGI1* overexpression enhances resistance to both CRR and FCR. When KO1 mutant plants and the Bobwhite control were challenged with CRR, both were highly susceptible, and no significant difference was observed (Fig. S13).

**Figure 6.**
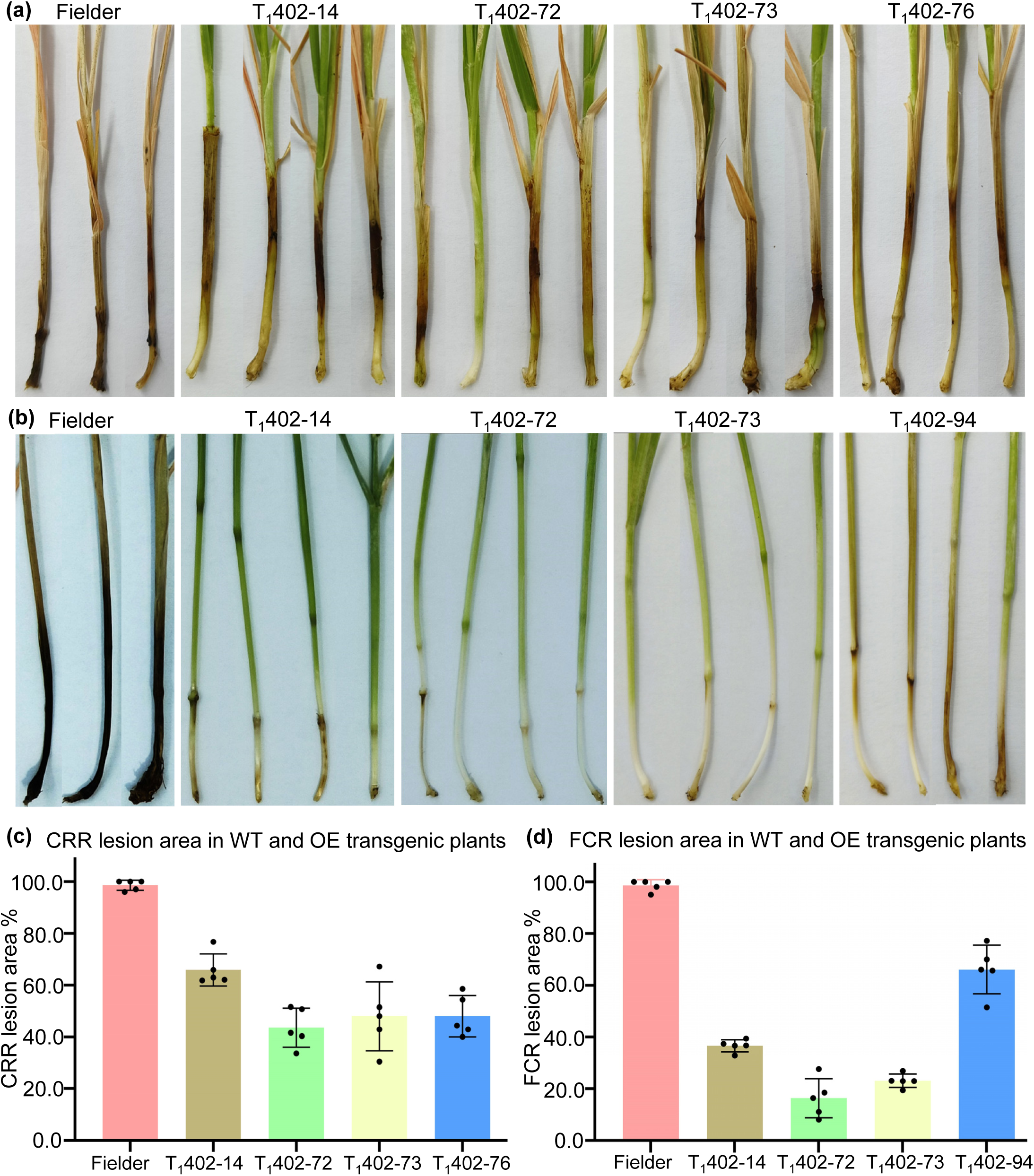
Overexpression of *BGI1* enhances resistance to common root rot (CRR) and *Fusarium* crown rot (FCR). (a-b) Phenotypic comparison of Fielder control and four OE transgenic families (T_1_402-14, T_1_402-72, T_1_402-73, and T_1_402-76/94) in response to CRR (a) and FCR (b). Plants were grown in growth chambers at 25℃ with a 16 h light / 8 h dark photoperiod. (c-d) Comparison of lesion areas in Fielder control and four OE transgenic families (T_1_402-14, T_1_402-72, T_1_402-73, and T_1_402-76/94) in response to CRR (c) and FCR (d). Error bars are standard errors of the means (n = 5). Lesion areas were determined from images of infected plants using ASSES v2 image analysis software.

**Figure 7.**
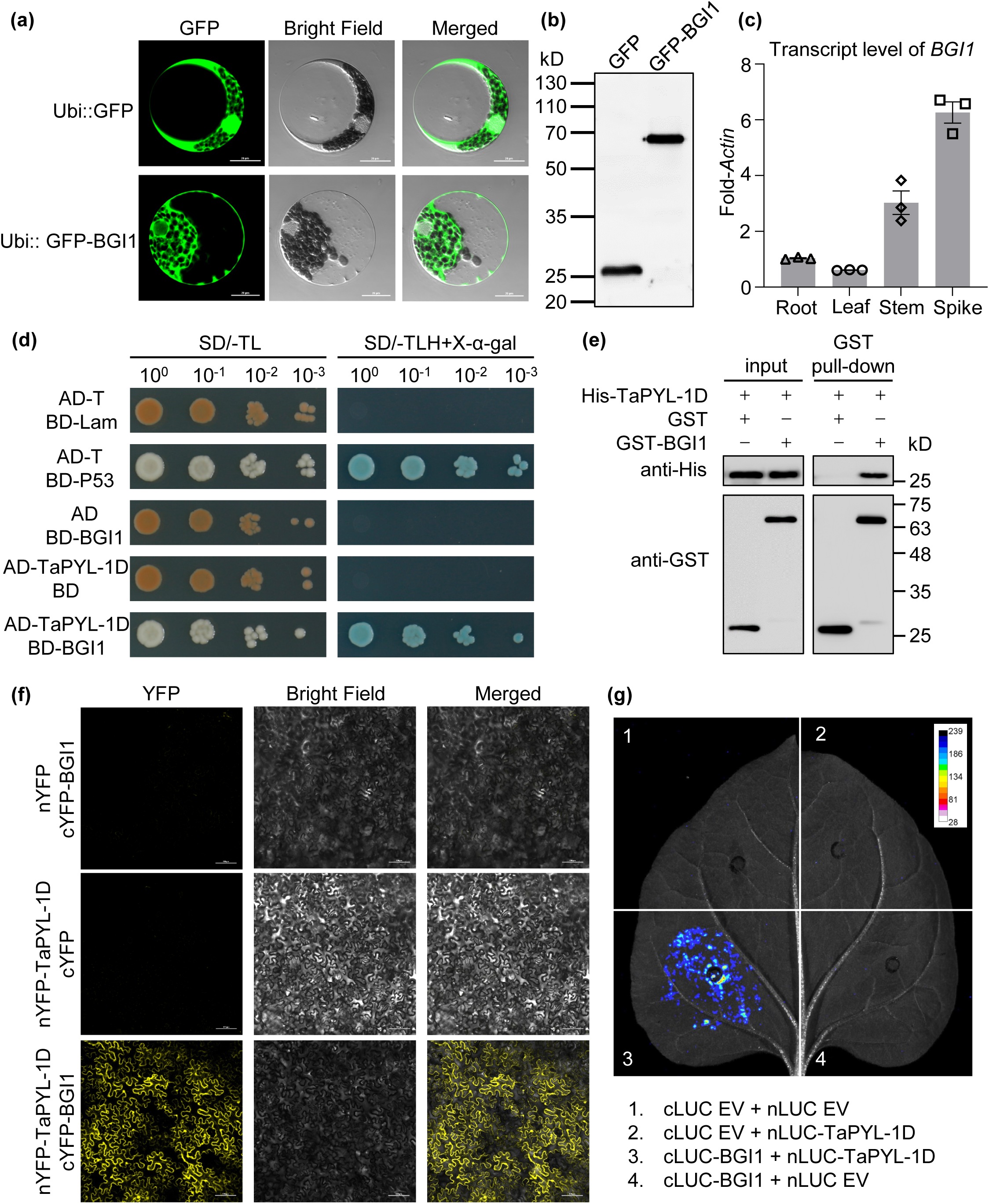
Characterization of BGI1. (a) Subcellular localization of Ubi::GFP and Ubi::GFP-BGI1 in wheat protoplasts. GFP, green fluorescent protein. Scale bars, 20 μm. (b) Western blot analysis of GFP and GFP-BGI1 proteins isolated from wheat protoplasts using an anti-GFP antibody. (c) Transcript levels of *BGI1* in different tissues of *Ae. tauschii* accession PI 511383. Transcript levels were quantified relative to *Actin* using the 2^-ΔCT^ method. Error bars indicate standard errors of the means (n = 3). (d) Yeast two-hybrid assays showing that BGI1 interacts with TaPYL-1D. AD, GAL4 activation domain; BD, GAL4DNA-binding domain; SD/-TL, synthetic dextrose medium lacking Trp and Leu; SD/-TLH + X-α-gal, synthetic dextrose medium lacking Trp, Leu, and His (X-α-gal = 40 mg·mL^D1^). (e) GST pull-down assays showing the interaction between BGI1 and TaPYL-1D *in vitro*. GST-BGI1 was incubated with His-TaPYL-1D, pulled down using anti-GST agarose beads, and detected by anti-His immunoblotting. (f) BiFC assays confirming the interaction between BGI1 and TaPYL-1D in tobacco epidermal cells. YFP, yellow fluorescent protein. Scale bars, 100 μm. (g) LCI assays confirming the interaction between BGI1 and TaPYL-1D in tobacco leaves. Scale bar represents luminescence intensity.

To further investigate the underlying mechanism of enhanced resistance, we analyzed the transcript levels of three pathogenesis-related (*PR*) genes in the stems of OE transgenic plants and Fielder at the heading stage. qRT-PCR experiments revealed that the transcript levels *PR1*, *PR4*, and *PR5* showed no significant difference between OE transgenic plants and the Fielder control (Fig. S14). These findings suggest that the enhanced resistance to pathogen infection in OE transgenic plants may be attributed to increased lignin content and the associated mechanical barrier, rather than induced systemic resistance.

### BGI1 has CAD activity

To characterize the enzymatic activity of BGI1, a fusion protein of glutathione S-transferase (GST)-BGI1 was expressed and purified in *Escherichia coli* expression system. Western blot analysis confirmed the expression of the recombinant GST-BGI1 protein with an expected molecular weight of 66 kD (Fig. 5j). The enzymatic activity of the purified recombinant GST-BGI1 protein was evaluated using cinnamaldehydes and NADPH as substrates, with the activity quantified by the rate of NADPH consumption (Chabannes et al., 2001). The results showed a statistically significant correlation between BGI1’s enzymatic activity and the concentration of the recombinant protein provided (Fig. 5k), demonstrating the in *vitro* CAD activity of the recombinant BGI1 protein.

To investigate the catalytic activity of native BGI1 *in vivo*, the KO mutant plants were further examined. A comparison of CAD activity in the internodes between WT Bobwhite and the KO1 mutant plants at the heading stage revealed a significant reduction in enzyme activity in KO mutant plants compared to WT (*P* < 0.001; Fig. 5I). Conversely, the CAD activities of OE transgenic lines (T_1_402-72 and T_1_402-73) were significantly increased compared to Fielder (Fig. S15). These findings demonstrate that BGI1 exhibits CAD activity both *in vitro* and *in vivo*.

### Subcellular localization and expression pattern of *BGI1*

To determine the subcellular localization of BGI1, we performed transient expression assays in wheat protoplasts using green fluorescent protein (GFP) and GFP-BGI1 fusion protein constructs derived by the UBI promoter. In contrast to the ubiquitous distribution of free GFP, the fluorescence signal of GFP-BGI1 was exclusively confined to the cytoplasm (Fig. 7a). Anti-GFP immunoblots confirmed that both GFP and GFP-BGI1 proteins were expressed at their expected sizes (Fig. 7b). Furthermore, quantitative reverse transcription PCR (qRT-PCR) analysis revealed that *BGI1* was expressed in all examined tissues, including the root, leaf, stem, and spike, with notably higher expression levels observed in the stem and spike (Fig. 7c).

### The regulatory networks and interaction protein of BGI1

To understand the regulatory networks associated with *BGI1*, we performed RNA-seq analysis to compare the transcriptomes of WT and KO mutants at the heading stage. This analysis identified 6,551 differentially expressed genes (DEGs) between WT and KO1 mutant plants (FDR < 0.01, |log_2_ fold change| > 1, *p* value < 0.05; Fig. S16a, Table S5). Among these, 2,345 DEGs were upregulated, while 4,206 were downregulated in the KO1 mutant plants compared to WT (Fig. S16a, Table S5). Gene Ontology (GO) annotations of these DEGs revealed significant enrichment in biological processes such as lignin biosynthetic process, pigment biosynthetic process, and abscisic acid (ABA) metabolic process (Fig. S16b). The RNA-seq data indicated that lignin biosynthesis and ABA-related genes were downregulated in KO1 plants compared to WT (Fig. S17). qRT-PCR analysis confirmed the downregulation of two lignin biosynthesis related genes *TaCOMT1* (*TraesCS3A02G534900*) and *TaLAC4* (*TraesCS3B02G392700*) in KO1 plants (Fig. S18). Conversely, these genes were upregulated in the OE transgenic lines compared to WT (Fig. S19). Furthermore, we validated the expression levels of ABA-related genes *TaABA1* (*TraesCS2A02G317000*, *TraesCS2B02G335400*, and *TraesCS2D02G314900*), *TaPP2C* (*TraesCS1B02G242300*), and *TaSnRK* (*TraesCS1B02G347100* and *TraesCS3B02G225800*), and found that they were downregulated in KO1 (Fig. S20), supporting the RNA-seq results.

To identify potential protein interactions with BGI1, a yeast two-hybrid (Y2H) screen was conducted using the coding sequence of *BGI1* as bait against a wheat cDNA library. The Y2H analysis successfully identified an interaction between BGI1 and TaPYL-1D (TraesCS1D02G126900), a protein with a PYR/PYL/RCAR-like domain known for its role in the ABA signaling pathway (Liu et al., 2021a). This interaction was further verified by GST pull-down, bimolecular florescence complementation (BiFC), and luciferase complementation imaging (LCI) assays (Figs. 7d-g). These findings suggest a potential regulatory mechanism involving BGI1 in lignification and the ABA signaling pathway.

## Discussion

### The *bgi1* mutant is defective in lignin biosynthesis

*CAD* mutants have been previously characterized in several plant species, including *bm1* in maize (Halpin et al., 1998), *bmr6* in sorghum (Li et al., 2015; Scully et al., 2016), *gh2* in rice (Zhang et al., 2006), and *Bdcad1* in *B. distachyon* (Bouvier d’Yvoire et al., 2013). While these studies identified *CAD* mutants, to our knowledge, no corresponding *CAD* mutants has been identified and characterized in wheat. Using map-based cloning and SNP analysis, we identified the *BGI1* gene encoding a CAD protein and found that *bgi1* is a lignin-deficient mutant, leading to a reddish-brown coloration phenotype in lignified tissues.

In maize and sorghum, the *bm1* and *bmr6* mutant plants exhibit reddish-brown pigments in the stalk pith and leaf midrib (Halpin et al., 1998; Saballos et al., 2009). In rice, the *gh2* mutant plants display pigments in the internode, hull, and basal leaf sheath (Zhang et al., 2006). Consistent with previous studies, our research identified the *bgi1* mutant plants show a reddish-brown coloration in the internode, spike, spike rachilla, glume, and lemma (Fig. 1). All the above *CAD* mutants exhibit reddish-brown coloration in lignified tissues, suggesting that these CADs play a conserved role in lignin biosynthesis across maize, sorghum, rice, and wheat. However, phenotype differences were observed, such as the reddish-brown coloration of the inner glumes in the *bgi1* mutant, a trait not seen in other mutants. The observed phenotypic variations could be attributed to two key factors. First, these CAD orthologous genes may have some degree of functional differentiation among different plant species. The maize *BM1* (Halpin et al., 1998) and sorghum *BMR6* (Li et al., 2015) genes were expressed strongly in leaves, while rice *GH2* (Zhang et al., 2006), *B. distachyon BdCAD1* (Bouvier d’Yvoire et al., 2013), and wheat *BGI1* genes were expressed slightly in the same tissue. Second, the type of mutation (missense or splice acceptor) may lead to different levels of reduction in CAD enzymatic activity. The *gh2* mutant has an amino acid (G185D) change in the causal gene (Zhang et al., 2006). In *Brachypodium*, the Bd4179 and Bd7591 mutant lines carry one (G192D) or two (G99V and S286F) amino acid changes in BdCAD1 (Bouvier d’Yvoire et al., 2013). In contrast, the *bgi1* mutant plants carry a mutation (G > A) in the splicing acceptor site of the first intron of *BGI1*, leading to premature termination (Fig. 2f). These different types of mutations may contribute to the phenotypic differences observed across various plant species.

The KO mutant plants of *BGI1* exhibited reduced CAD activity and lower lignin content (Figs. 5 and S11), which is consistent with observations in some monocotyledonous plants, such as maize (Halpin et al., 1998; Xiong et al., 2020), sorghum (Scully et al., 2016), rice (Zhang et al., 2006), and *B. distachyon* (Bouvier d’Yvoire et al., 2013). However, estimates of lignin concentration can vary greatly depending on the extraction method used. Some studies have observed reduced CAD activity without a corresponding decrease in lignin content (Baucher et al., 1999; Baucher et al., 1996; Christiane Marque et al., 1998).

### Overexpressing *BGI1* results in increased lignin content and shearing force

Numerous studies have documented *CAD* mutants, yet research on the phenotypic changes resulting from *CAD* overexpression in cereal crops remains relatively scarce. In *Artemisia annua*, overexpression of the *AaCAD* gene led to significantly higher lignin content in transgenics compared with WT plants (Ma et al., 2018). Transgenic *Arabidopsis* plants overexpressing *IbCAD1* exhibited increased lignin content in stems and roots, with a higher proportion of S lignin compared to G lignin (Kim and Huh, 2019). *PpCAD2* overexpression transgenic tomato plants contained a higher lignin content and CAD enzymatic activity in the leaf, stem, and fruit pericarp tissues (Li et al., 2019). Although none of these studies were conducted in cereal crops, these results support our observation that overexpression of *BGI1* leads to increased lignin content (Fig. S11).

In addition to increasing lignin content, *BGI1* overexpression significantly enhances stem strength, as evidenced by increased shearing force of stems (Fig. S12). Recent correlation analyses have identified a significant positive relationship between *TaCAD1* gene expression and stem strength (Chen et al., 2021). Studies on wheat culms have demonstrated that lodging is a common issue in wheat production, causing yield losses ranging from 10% to 80% (Easson et al., 1993; Peng et al., 2014). Lignin accumulation has been positively and significantly correlated with internode breaking strength and culm lodging resistance (Peng et al., 2014). Thus, overexpression of *BGI1* could be a potential strategy to improve lodging resistance in wheat.

Interestingly, overexpression of *PpCAD2* in tomato resulted in increased plant height, longer roots, and larger stem diameter (Li et al., 2019). Overexpression of *IbCAD1* in *Arabidopsis* enhanced seed germination rate and increased tolerance to reactive oxygen species (Kim and Huh, 2019). Although our *BGI1* overexpression transgenic plants did not appear to show significant phenotypic changes (Fig. S21), comprehensive agronomic and quality evaluations are necessary.

### Overexpression of *BGI1* enhances pathogen resistance

Yield losses caused by *B. sorokiniana* are often severe, ranging from 15% to 20%. Under favorable conditions of heat and drought, this disease can reduce wheat production by up to 70% and cause significantly seed quality deterioration (Sharma and Duveiller, 2007). FCR infection is estimated to cause a 35% yield loss in winter wheat in the Pacific Northwest region of the United States, with wheat seeds likely to become contaminated with fungal toxins (Smiley et al., 2005; Su et al., 2021). Additionally, there are only a limited number of genetic loci conferring resistance to CRR or FCR (Su et al., 2021). In our study, transgenic plants overexpressing *BGI1* exhibited increased resistance against both CRR and FCR infections (Figs. 6a, b). This enhanced resistance to pathogen infections in OE transgenic plants is linked to the increased lignin content.

Lignin serves as a mechanical defense barrier and is known to be involved in plant defense against pathogen infections (Miedes et al., 2014; Vance et al., 1981). Lignification can form protective barriers against pathogen invasion, modify cell walls to resist pathogen-released degrading enzymes, enhance the resistance of cell walls to toxins diffusion from pathogens to hosts, generate free radicals and toxic precursors, and lignify to entrap pathogens (Bhuiyan et al., 2009; Rong et al., 2016). In *Arabidopsis*, genetic and functional analyses indicate that *CAD-C* and *CAD-D* not only serve as key enzymes in lignin biosynthesis but also play vital roles in plant defense against *Pseudomonas syringae* pv. *tomato* (Tronchet et al., 2010). In wheat, increased lignin accumulation has been associated with increased resistance against FCR (Yang et al., 2021). Likewise, wheat lines overexpressing *TaCAD12* demonstrated significantly enhanced resistance to the fungus *Rhizoctonia cerealis* throughout all growth stages (Rong et al., 2016). These results support our findings that overexpression of *BGI1* leads to increased pathogen resistance.

In *Arabidopsis*, *cad-C* and *cad-D* mutations negatively influenced the expression of *PR1* and *PR5* genes after inoculation with *Pseudomonas syringae* pv. *Tomato* (Tronchet et al., 2010). In wheat, transcriptional levels of *PR10* and *PR17*c were significantly elevated in the stems of *TaCAD12*-overexpression wheat plants, contributing to increased resistance against *R. cerealis* (Rong et al., 2016). However, our study did not observe changes in transcript levels of *PR* genes in *BGI1*-overexpressing plants upon pathogen inoculation (Fig. S14), suggesting that *CAD*-overexpression may involve different mechanisms in plant defense. The precise molecular mechanisms underlying the pathogen resistance caused by *BGI1* overexpression require further investigation.

### The interaction between BGI1 and TaPYL-1D may contribute to the process of lignification

Previous studies have reported that the hormone ABA plays a crucial role in regulating plant secondary cell-wall deposition and lignification (Brookbank et al., 2021; Liu et al., 2021a). ABA signaling is mediated by intracellular PYL receptors, which bind to and inhibit PP2Cs, thereby releasing protein kinases SnRK2s from inhibition (Chen et al., 2020; Park et al., 2009). In muskmelon, exogenous ABA enhanced CAD activity by 20.47% and promoted the production of lignin monomers, lignin content, and phenolic acids in fruit wounds (Wang et al., 2024b). In Kenaf (*Hibiscus cannabinus* L.), ABA treatment induced a biphasic expression pattern of *HcCAD2*, with significant induction at 6 h, 24 h, and 48 h (Choi et al., 2016). Additional studies revealed that ABA increased the expressions of lignin biosynthesis genes, such as *CAD*, *4-coumarate-CoA ligase*, and *cinnamate 4-hydroxylase*, as well as the activities of lignifying enzymes, thereby promoting lignin accumulation (Cheng et al., 2013; Liu et al., 2021b; Xu et al., 2020). These studies suggest a potential regulatory mechanism involving ABA, PYL receptors, and CAD proteins in lignin biosynthesis. Our study identified a physical interaction between BGI1 and TaPYL-1D using Y2H, LCI, BiFC, and GST pull-down assays (Figs. 7d-g). This finding aligns with GO annotations indicating enrichment in the ABA metabolic process (Fig. S16b), suggesting a regulatory role for BGI1 in ABA signaling. Our results provide a foundation for further investigation into the functional mechanisms of BGI1 and TaPYL-1D in regulating lignin biosynthesis in wheat.

In summary, we successfully isolated the *bgi1* gene in wheat, which encodes the TaCAD1 protein. Functional validation through loss-of-function EMS and CRISPR mutations in *BGI1* result in a reddish-brown coloration of lignified tissues. Disruption of *BGI1* significantly reduces lignin content without affecting plant development, potentially enhancing the use of wheat residues for biofuel production and other bio-based products. Conversely, overexpression of *BGI1* elevates lignin content and boosts pathogen resistance, which could lead to the development of wheat varieties with improved disease tolerance and lodging resistance. Specifically, increased disease resistance for CRR and FCR could reduce reliance on chemical control methods, contributing to more sustainable agricultural practices.

## Materials and Methods

### Plant materials and mapping populations

Mutant lines were generated by treating approximately 14,800 seeds from the *Ae. tauschii* accession PI 511383 with 0.4% EMS (Sigma-Aldrich Co., MO, USA). Seeds from 2,156 independent M_1_ mutant plants were harvested. We identified an M_2_ mutant line (M326) segregating plants with reddish-brown pigmentation in various tissues. The M_2_ plants exhibiting this pigmentation were transplanted and self-pollinated to produce a homozygous M_3_ family. As the M_2_ seeds of M326 were exhausted, the M_3_ plants were crossed with the WT PI 511383, generating an F_2_ population of 1,872 individuals. To accelerate the development of molecular markers, we generated an additional mapping population from a cross between M326 and AL8/78 (Luo et al., 2017), a genotype which is highly polymorphic compared to PI 511383.

### BSR-Seq analysis

Based on the phenotypic data, 20 F_2_ plants displaying the WT phenotype and 20 F_2_ plants exhibiting the mutant phenotype from the M326 × PI 511383 cross were selected to generate the wild bulk (W-bulk) and mutant bulk (M-bulk) for BSR-Seq analysis. Total RNA was isolated from both parents and W/M-bulks using the NucleoZol reagent (Macherey-Nagel, Düren, Germany) and purified with the Direct-zol RNA MiniPrep Plus Kit (Zymo Research, CA, USA). RNA sequencing was carried out at Novogene Bioinformatics Technology Co., Ltd. (Beijing, China). The raw sequencing data is available in the National Genomics Data Center (NGDC) database under the accession number PRJCA028685. Procedures for sequencing reads analysis and variant calling were adopted from previously reported methods (Bai et al., 2024; Jiang et al., 2023). SNP-index and Δ|SNP-index| were calculated across the genome to identify potential genomic regions of interest (Takagi et al., 2013).

### Development of CAPS and KASP markers

SNPs identified within the candidate region on chromosome 6D were used to develop both CAPS (Konieczny and Ausubel, 1993) and KASP markers (Semagn et al., 2014). For CAPS markers, primers were designed using Primer3 (https://bioinfo.ut.ee/primer3-0.4.0/primer3/). The resulting PCR products were digested with appropriate restriction endonucleases (New England BioLabs Inc., Hitchin, UK). For KAPS markers, forward primers were designed with standard FAM-compatible tails (5′-GAAGGTGACCAAGTTCATGCT-3′) or HEX tails (5′-GAAGGTCGGAGTCAACGGATT-3′), incorporating the polymorphism between the parents at the 3′ end. Common reverse primers were chromosome 6D.

### Phylogenetic analysis

specifically designed for Using the sequence of *BGI1* as a query, homeologous sequences from various plant species were obtained from the publicly available plant genome database (http://plants.ensembl.org/). Protein sequences were aligned using Muscle, as implemented in the software MEGA v7 (Kumar et al., 2016). Phylogenetic trees were constructed using the Neighbor-Joining method. The resulting phylogenetic tree was visualized using the Interactive Tree Of Life (iTOL) v6 (https://itol.embl.de/).

### EMS mutants in tetraploid and hexaploid wheat

The sequenced EMS mutagenized populations of the tetraploid wheat variety Kronos and the hexaploid cultivar Cadenza (Krasileva et al., 2017) are available on line at https://www.wheat-tilling.com/. Using BLASTN with the sequence of *BGI1* as a query, we identified two Kronos mutant lines (K3288 and K3088) that harbor premature stop codons in *BGI-A1* and *BGI-B1*, respectively. We crossed the K3288 mutant with the K3088 mutant, self-pollinated the F_1_ plants, and selected F_2_ plants that were homozygous for *BGI1* homeologs-WT, *bgi-A1*, *bgi-B1*, and the *bgi1* double mutant.

Similarity, we identified three Cadenza mutant lines (Cadenza0376, Cadenza0749, and Cadenza0183) carrying premature stop codons in *BGI-A1*, *BGI-B1*, and *BGI-D1*, respectively. Through crossing and selection in the F_2_ generation, we generated single-gene mutants (*bgi-A1*, *bgi-B1*, and *bgi-D1*), double mutants (*bgi-A1B1*, *bgi-A1D1*, and *bgi-B1D1*), and the *bgi1* triple mutant (*bgi-A1B1D1*).

### BSMV-mediated gene editing and OE transgenic validation

The previously reported plasmids used for BSMV-mediated gene editing in wheat were employed (Li et al., 2021). The resulting BSMV-sgRNA constructs were transformed into *A. tumefaciens* strain EHA105 and subsequently co-infiltrated into *N. benthamiana* leaves. The inoculated leaves were then homogenized and rub-inoculated onto the leaves of Bobwhite *Cas9*-transgenic plants. Sanger sequencing was used to identify plants homozygous for single-gene, double, and triple mutants in the M_2_ population.

The coding sequence of *BGI1* was introduced into the modified binary vector pCAMBIA1300 and then transformed into the hexaploid wheat variety Fielder via *A. tumefaciens*-mediated transformation at the Peking University Institute of Advanced Agricultural Sciences transformation facility.

### Subcellular localization assay

The coding sequence of *BGI1* was cloned into the pJIT163-Ubi-GFP vector (Luo et al., 2022). Both the empty vector (EV) and the resulting *ubi*::GFP-BGI1 construct were transformed into wheat protoplasts (variety Fielder) using the PEG-mediated method (Luo et al., 2022). Protoplast images were captured using a confocal microscope (A1 HD25 Nikon, Tokyo, Japan).

### qRT-PCR and transcriptome analysis

Stem, leaf, and spike samples from *Ae. tauschii* accession PI 511383 were collected at the heading stage, while root samples were collected at the three-leaf stage. qRT-PCR was conducted on an Applied Biosystems ABI QuantStudio 5 Real-Time PCR System (Applied Biosystems, CA, USA). RNA expression levels were normalized using the endogenous control *TaActin* (Zhang et al., 2017), and *BGI1* transcript levels were calculated using the 2^-ΔCT^ method as previously described (Chen et al., 2018). The primers for *PR* genes were previously described (Zhang et al., 2017).

Differentially expressed genes (DEGs) were identified using edgeR software (Robinson et al., 2010). Genes showing a false discovery rate (FDR) < 0.01, |log_2_ fold change| > 1, and *p*-value < 0.05 were classified as DEGs. GO enrichment analysis of DEGs was conducted using agriGO v2.0 (http://systemsbiology.cau.edu.cn/agriGOv2/).

### BGI1 protein expression and purification in *E. coli*

The coding sequence of *BGI1* was inserted into the pGEX-4T-1 vector using the In-Fusion HD cloning kit (Takara Biotech, Shiga, Japan). Subsequently, the pGEX-4T-1 and GST-BGI1 constructs were transformed into the *E. coli* expression strain BL21. Protein expression was achieved by adding 0.3 mM isopropyl-β-D-thiogalactopyranoside (IPTG) for 16 hours at 16℃. Purification of the GST and recombinant GST-BGI1 proteins was carried out using Glutathione Magnetic Agarose Beads (Thermo Fisher, Waltham, MA, USA). The purified proteins were quantified using a BCA protein assay kit (Cwbio, Jiangsu, China).

### Western blotting analysis

Purified proteins were separated using sodium dodecyl sulfate-polyacrylamide gel electrophoresis (SDS-PAGE) (GenScript, Nanjing, China). The separated proteins were then electrotransferred onto a polyvinylidene fluoride (PVDF) membrane (Roche, Basel, Switzerland). Immunoblotting was conducted using a commercial HRP-conjugated Mouse anti GST-Tag antibody (Abclonal, Wuhan, China). A primary anti-GFP antibody (Abcam, Cambridge, UK) and a secondary Goat anti-Rabbit IgG-HRP antibody (Abmart, Shanghai, China) were used in subcellular localization assays.

### BGI1 enzyme activity assay

Enzyme activity assays were conducted on second internode tissues (∼100 mg) following the methods described previously (Chabannes et al., 2001). Both *in vivo* and *in vitro* CAD activities were measured using the CAD Activity Assay Kit (Sangon Biotech, Shanghai, China).

### Histochemical staining

Wiesner staining (Blaschek et al., 2020) and Mäule staining (Sibout et al., 2005) were used to detect the localization of lignin accumulation. For Wiesner staining, fresh hand-cut cross-sections of wheat second internodes at the heading stage were initially incubated in a phloroglucinol solution for 10 minutes, followed by a 5-minute treatment with 18% HCl. For Mäule staining, fresh cross-sections were sequentially treated: 1% KMnO_4_ for 15 minutes, rinsed with water, 30% HCl for 1 minute, rinsed again, and then mounted in 5% NaHCO_3_ for 10 minutes (Sibout et al., 2005). After treatment, the cross-sections were immediately photographed using a Leica microscope (Leica, Hamburg, Germany).

### Lignin content and composition analysis

Internodes were collected at the heading stage, oven-dried, and then ground into powder. The power was sieved through a 40-mesh screen for lignin content and composition analysis. Klason lignin was determined using the previously reported method (Zhang et al., 2021).

### Y2H assays

A yeast two-hybrid assay was conducted following the Clontech Yeast Two-Hybrid System protocol (Clontech, Mountain View, CA, USA). BGI1 was used as bait for screening a cDNA library constructed from Fielder (Li et al., 2023). To validate the interaction between BGI1 and TaPYL-1D, the *BGI1* coding sequence was ligated into the pGBKT7 vector, while the *TaPYL-1D* coding sequence was cloned into the pGADT7 vector. The constructs were co-transformed into the yeast strain Y2H Gold and cultured on SD/-Trp/-Leu medium. Co-transformants were then plated on selective medium (SD/-Trp/-Leu/-His + 40 μg/ml X-α-gal).

### *In vitro* GST pull-down assays

The *BGI1* coding sequence was inserted into the pGEX-4T-1 vector to generate a GST-tagged BGI1 protein. Similarly, the *TaPYL-1D* coding sequence was cloned into the pET-28a vector to produce His-tagged TaPYL-1D protein. Both GST and His fusion proteins were purified using glutathione magnetic agarose beads and HisPur Ni-NTA magnetic beads (Pierce, Waltham, MA, USA), respectively. Pull-down assays were performed following the manufacturer’s protocol (Pierce, Waltham, MA, USA). The bait-prey mixtures were denatured at 95℃ for 10 minutes, separated by 10% SDS-PAGE, and immunoblotted with anti-His (Abclonal, Wuhan, China) and anti-GST antibodies (Abclonal, Wuhan, China).

### *In vivo* BiFC assays

The coding sequence of *BGI1* was inserted into pCAMBIA1300-35S-cYFP, while TaPYL-1D was fused with pCAMBIA1300-35S-nYFP. These constructs were transformed into *Agrobacterium* strain EHA105 and then infiltrated into 4-week-old *N. benthamiana* leaves. YFP fluorescence signals were detected two days post-infiltration using a confocal microscope (A1 HD25 Nikon, Tokyo, Japan).

### LCI assays

The coding sequences of *BGI1* and *TaPYL-1D* were inserted into the pCAMBIA1300-cLuc and pCAMBIA1300-nLuc vectors, respectively (Zhou et al., 2018). The constructs, cLuc-BGI1 and nLuc-TaPYL-1D, were transformed into *Agrobacterium* strain EHA105 and co-infiltrated into 4-week-old *N. benthamiana* leaves. Luciferase activities were captured 48 hours post-infiltration using the ChemiDoc MP Imaging System (Bio-Rad, Hercules, USA). Fluorescence signals in the injected leaves were quantified using ImageJ software (Schneider et al., 2012).

### Evaluation of pathogen resistance

Inoculation experiments with *B. sorokiniana* and *Fusarium pseudograminearum* were performed at Hebei Agricultural University, Baoding, Hebei, China. The plants were grown in growth chambers under controlled conditions (25℃ and 16 h photoperiod). The dominant strains of *B. sorokiniana* and *F. pseudograminearum* (Wz2-8A) prevalent in Hebei Province, China, were isolated from infected wheat plants and cultured on potato dextrose agar (PDA) medium. Procedures for inoculation and statistical analyses of infection types have been previously reported (Wang et al., 2024a; Yuan et al., 2024).

## Author contributions

LH, RS, and XH performed most of the experimental tasks; JZ conducted the BGI1 enzymatic activity assays; YL contributed subcellular localization; JL contributed mapping; HL contributed expression analysis; GW designed the *BGI1* sgRNA; SR contributed to the writing process; JW and DF provided and analyzed EMS mutants; XW and XR designed and performed the pathogen infection experiments; CZ developed and screened the PI 511383 EMS mutant population. LH and SC analyzed the data and wrote the first version of the manuscript. SC, CZ, XW, and YD proposed and supervised the project. SC generated the final version of the paper. All authors revised the manuscript and provided suggestions.

## Acknowledgements

The authors thank Dr. Jorge Dubcovsky (University of California, Davis, USA) for providing the Kronos and Cadenza mutant lines. Work at SC laboratory was supported by the Key R&D Program of Shandong Province (ZR202211070163, 2023LZGC022), the Young Yuandu Scholars Program of Weifang City, and the Young Taishan Scholars Program of Shandong Province. Work at YD laboratory was supported by the Shandong Provincial Natural Science Foundation (ZR2020MC029). Work at XW laboratory was supported by the National Key Research and Development Program of China (2023YFD1201002), Provincial Natural Science Foundation of Hebei (C2022204010), and S&T Program of Hebei (23567601H).

## Data availability

Data supporting the findings of this work are available within the paper and its supplementary information files. All the raw sequencing data generated for this project are archived at the National Genomics Data Center under BioProject accession number PRJCA028685.

## Supporting information

Additional supporting information may be found online in the Supporting Information section at the end of the article.

**Figure S1** Genetic map of *bgi1* based on 182 F_2_ plants from the M326 × AL8/78 cross and ten molecular markers.

**Figure S2** Expression levels of *AET6Gv20439300* (a) and *AET6Gv20438400* (b) homeologs.

**Figure S3** Phylogenetic analysis of BGI1.

**Figure S4** Sequence alignments of AET6Gv20438400 protein and five representative homeologs.

**Figure S5** Validation of *BGI1* using Cadenza EMS mutants.

**Figure S6** Procedures for the generation of single-gene mutants, double mutants, and the *bgi1* triple mutant.

**Figure S7** Generation of *BGI1* genome editing mutant plants using the BSMV-sg system.

**Figure S8** Phenotypic comparison between WT Bobwhite and mutant lines generated using BSMV-sgRNA-based gene editing approach.

**Figure S9** Statistical analysis of agronomic traits in Bobwhite and *BGI1* KO1 mutant plants.

**Figure S10** Transcript levels of *BGI1* in overexpression transgenic T_0_ plants.

**Figure S11** Total lignin content and Klason lignin levels in stems.

**Figure S12** Shearing force of WT, KO1 mutant plants, and OE transgenic lines (T_1_402-72 and T_1_402-73).

**Figure S13** Typical common root rot (CRR) symptoms and lesion areas in Bobwhite and KO1 mutant plants.

**Figure S14** Expression levels of pathogenesis-related (*PR*) genes in Fielder and *BGI1* overexpressing T_1_ plants.

**Figure S15** Comparison of CAD activities between Fielder and OE transgenic lines (T_1_402-72 and T_1_402-73).

**Figure S16** Transcriptome analysis between KO1 mutant plants and Bobwhite.

**Figure S17** Heatmap showing representative differentially expressed genes involved in lignin biosynthesis and ABA-related pathways.

**Figure S18** Expression levels of lignin biosynthesis-related genes in Bobwhite and KO1 mutant plants.

**Figure S19** Expression levels of lignin biosynthesis-related genes in Fielder and *BGI1* overexpressing T_1_ plants.

**Figure S20** Expression levels of ABA-related genes in KO1 mutant plants and Bobwhite.

**Figure S21** Phenotypic comparison of Fielder and three *BGI1* overexpression transgenic lines.

**Table S1** SNPs associated with the *bgi1* locus on *Aegilops tauschii* chromosome 6D.

**Table S2** Primers used in this study.

**Table S3** Predicted genes within the *bgi1* candidate region.

**Table S4** Genotypes of 99 M_1_ plants derived from the selected M_0_ edited plants.

**Table S5** Differentially expressed genes (DEGs) between Bobwhite and KO1 mutant plants.

