## Supplementary Figures for "Manipulation of the *brown glume and internode 1* gene leads to alterations in lignified tissue coloration, lignification, and pathogen resistance in wheat"

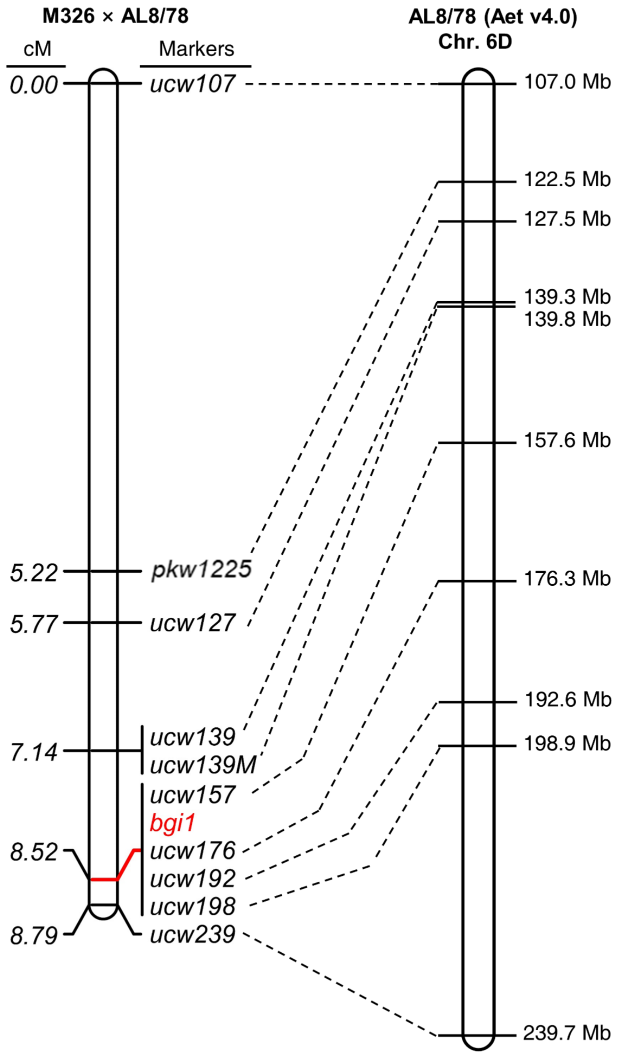


**Figure S1** Genetic map of *bgi1* based on 182 $F_2$ plants from the M326 × AL8/78 cross and ten molecular markers. Coordinates are based on the reference genome of the *Aegilops tauschii* accession AL8/78 (Aet v4.0). The values to the left of the PCR markers represent the genetic distances in centimorgans (cM). Mb, megabases.

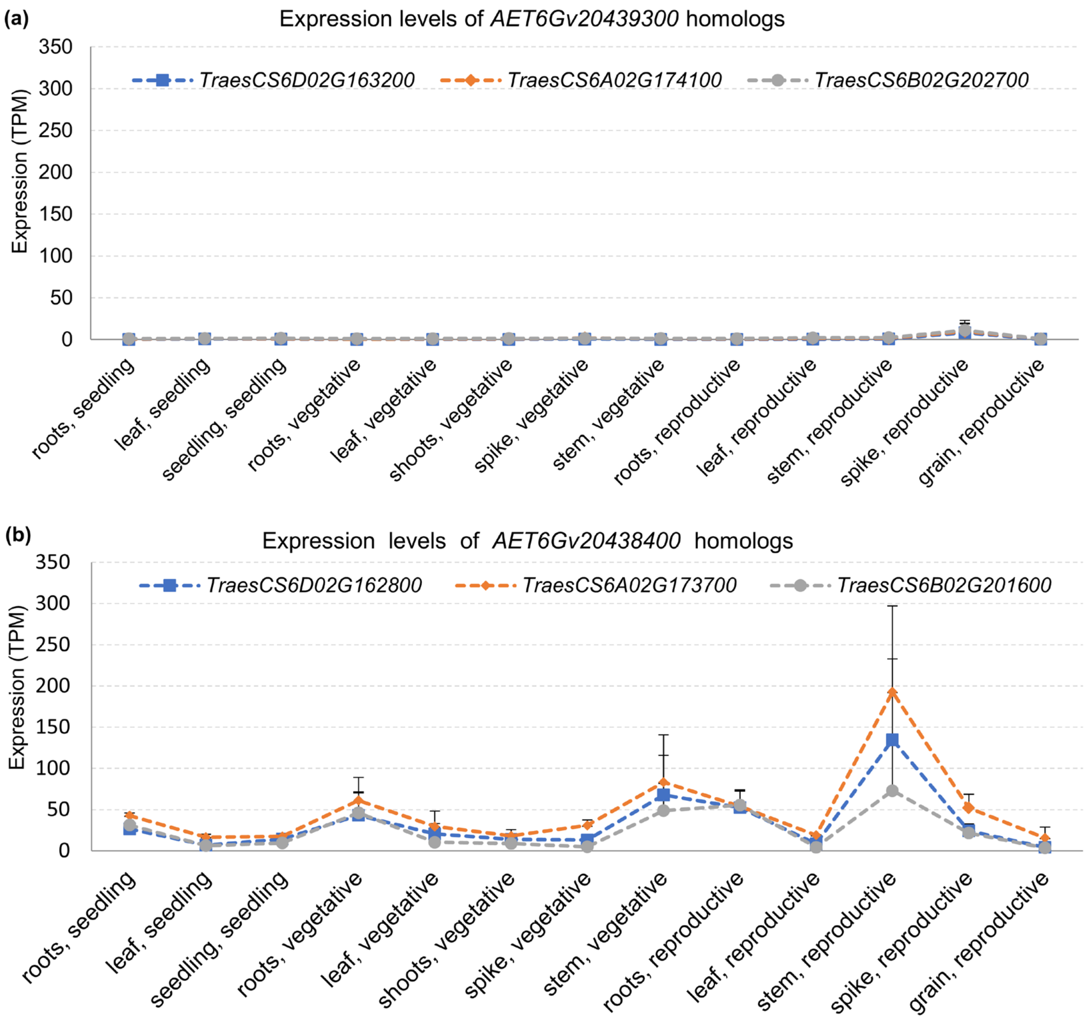


**Figure S2** Expression levels of *AET6Gv20439300* (a) and *AET6Gv20438400* (b) homeologs. RNA-seq data were published in the wheat expVIP database (http://www.wheat-expression.com/). TPM, transcripts per million; Error bars are standard errors of the means (n≥3).

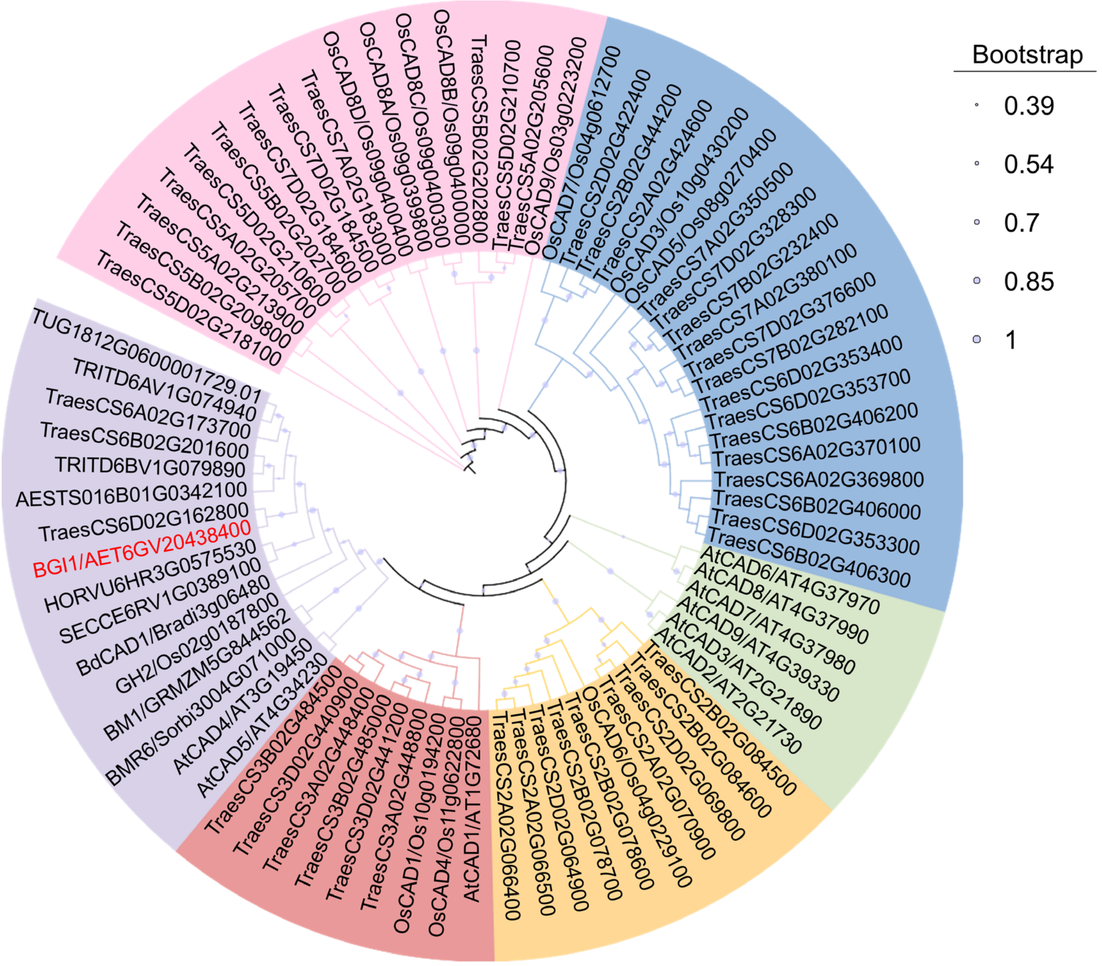


**Figure S3** Phylogenetic analysis of BGI1. Protein sequences were sourced from various plant species, including rice (Os), *Arabidopsis* (At), maize (GRMZM), *Sorghum bicolor* (Sorbi), *Aegilops tauschii* (AET), *Triticum urartu* (TU), *Aegilops speltoides* (AES), *Triticum turgidum* (TRITD), *Brachypodium distachyon* (Bradi), *Secale cereale* (SECCE) and *Hordeum vulgare* (HORVU). Protein sequences were aligned using Muscle, as implemented in the software MEGA v7. The phylogenetic tree was constructed using the Neighbor-Joining method (pair-wise deletion, bootstrap confidence values based on 1000 iterations). The resulting phylogenetic tree was visualized using the Interactive Tree Of Life (iTOL) v6 (https://itol.embl.de/). BGI1/AET6GV20438400 is highlighted in red. Two proteins, Ae.speltoides.TS01.6B01G0342100 and HORVU.MOREX.r3.6HG0575530, were abbreviated as AESTS016B01G0342100 and HORVU6HR3G0575530, respectively.

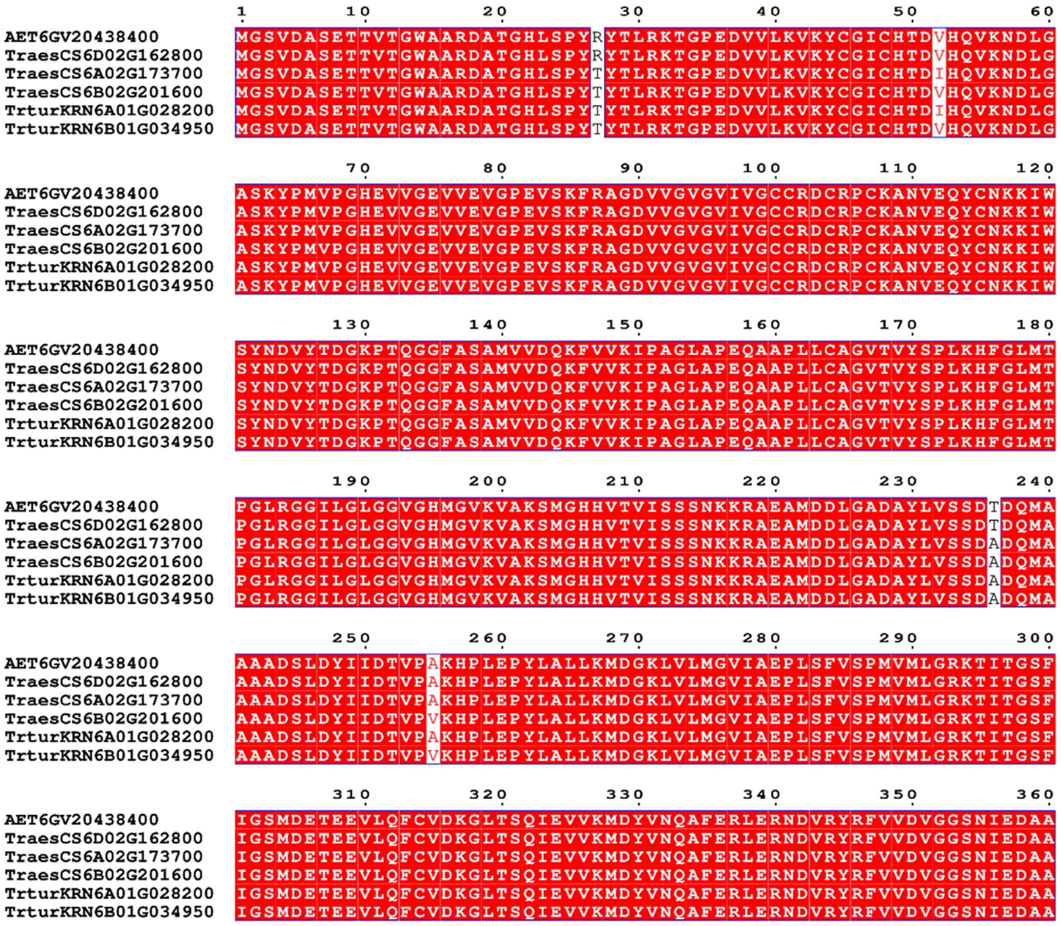


**Figure S4** Sequence alignments of AET6Gv20438400 protein and five representative homeologs. Protein sequences were sourced from the sequenced reference genomes of Chinese Spring (TraesCS) and Kronos (TrturKRN). Highlighted in red are identical sequences among these proteins. Sequences were aligned using the muscle function implemented in MEGA v7. ESPript 3.0 was used to visualize the alignments (https://espript.ibcp.fr/ESPript/cgi-bin/ESPript.cgi).

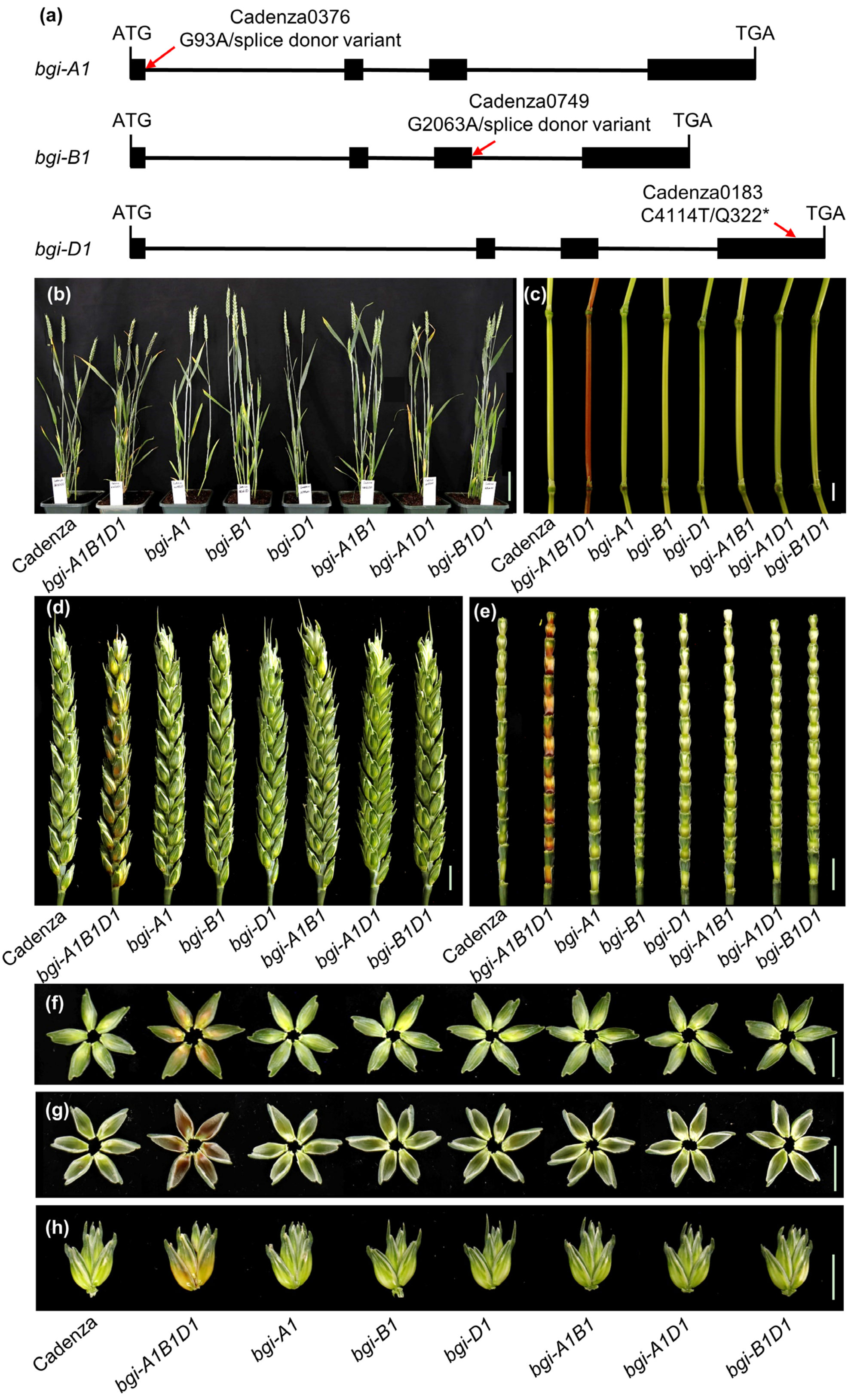


**Figure S5** Validation of *BGI1* using Cadenza EMS mutants. (a) Gene structure of the *BGI1* homeologs in the hexaploid wheat variety Cadenza. Exons are represented by black boxes and introns by black lines. EMS mutation sites are highlighted by red arrows. (b-i) Phenotypic comparison of the plant (b), internode (c), spike (d), spike rachilla (e), outer glume (f), inner glume (g), and spikelet with glumes removed (h) among Cadenza (WT), single-gene mutants (*bgi-A1*, *bgi-B1*, and *bgi-D1*), double mutants (*bgi-A1B1*, *bgi-A1D1*, and *bgi-B1D1*), and the triple mutant (*bgi-A1B1D1*) at the heading stage. Scale bars, 10 cm in (b) and 1 cm in (c-h).

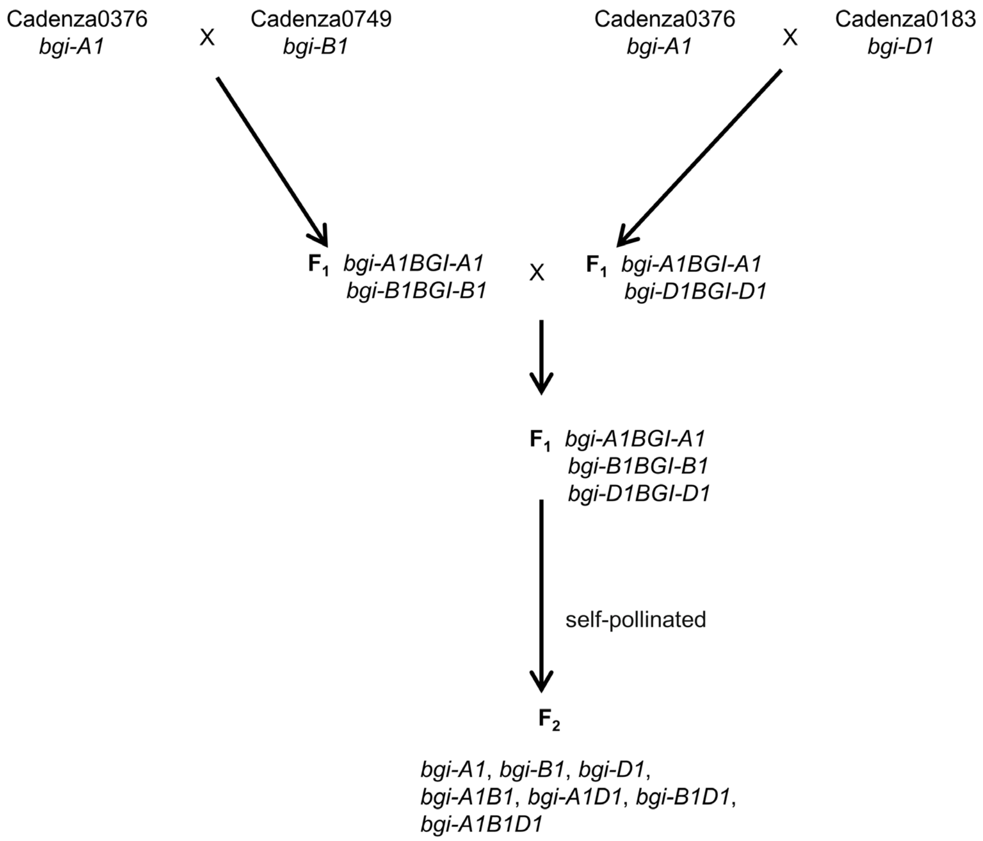


**Figure S6** Procedures for the generation of single-gene mutants, double mutants, and the *bgi1* triple mutant. The mutant lines Cadenza0376 (*bgi-A1*), Cadenza0749 (*bgi-B1*), and Cadenza0183 (*bgi-D1*) were selected from the sequenced EMS-mutagenized population of the hexaploid wheat variety Cadenza (https://www.wheat-tilling.com/).

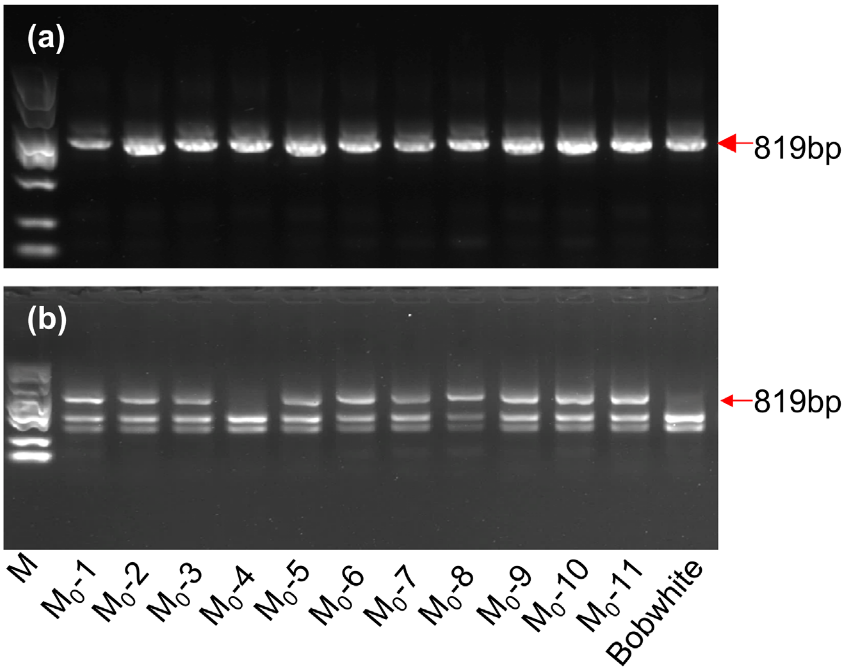


**Figure S7** Generation of *BGI1* genome editing mutant plants using the BSMV-sg system. (a) PCR products of the marker *HL553* (Table S2) from eleven infected $M_0$ wheat plants. (b) PCR products digested with *Psh*AI. The undigested 819-bp PCR products are highlighted by red arrows. $M_0$-1 to $M_0$-11 represent the infected $M_0$ plants in *Cas9*-transgenic Bobwhite; M, DNA ladder.

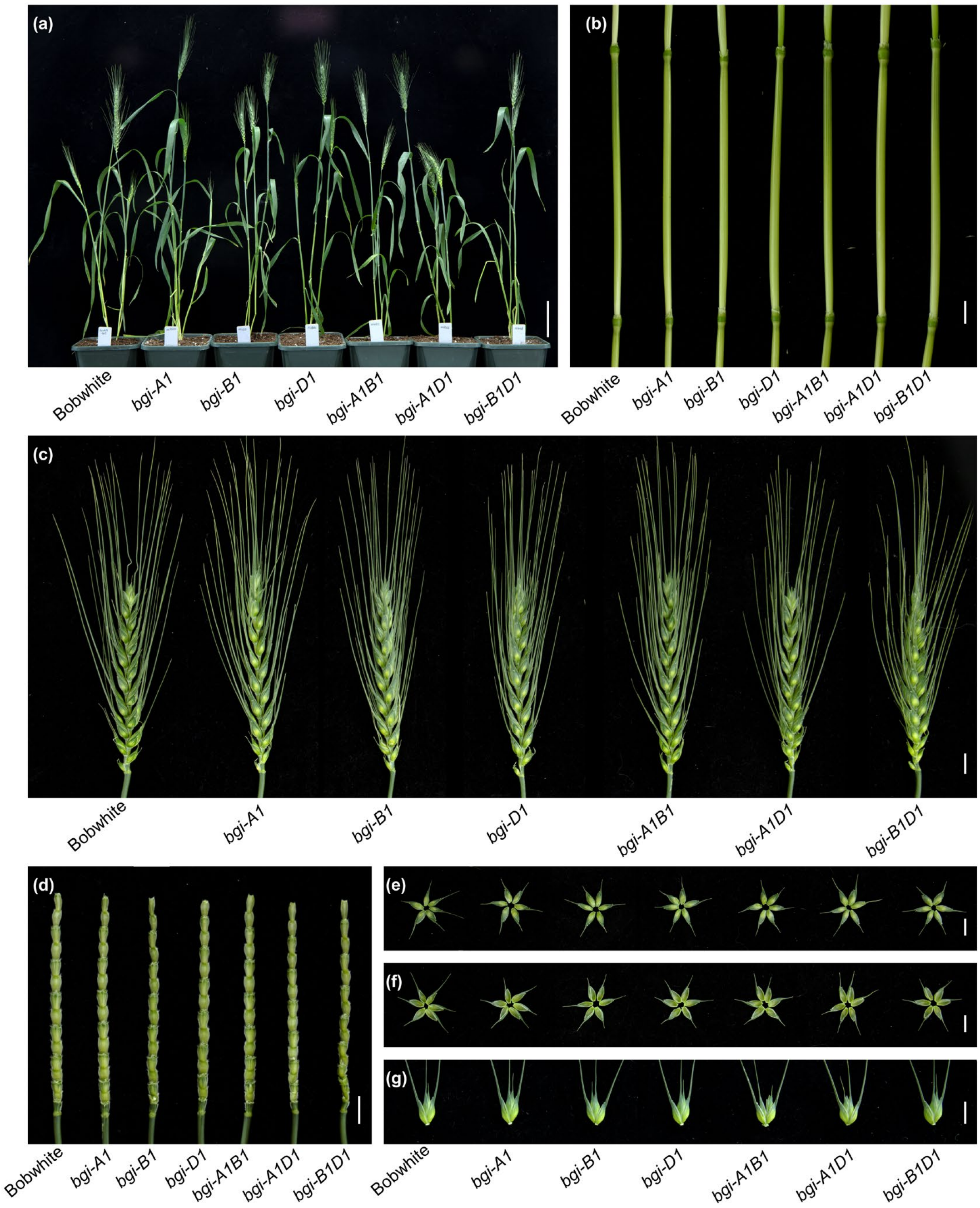


**Figure S8** Phenotypic comparison between WT Bobwhite and mutant lines generated using BSMV-sgRNA-based gene editing approach. (a-g) Phenotypic comparison of the plant (a), internode (b), spike (c), spike rachilla (d), outer glume (e), inner glume (f), and spikelet with glumes removed (g) between Bobwhite and the *bgi1* mutant lines (*bgi-A1*, *bgi-B1*, *bgi-D1*, *bgi-A1B1*, *bgi-A1D1*, and *bgi-B1D1*) at the heading stage. Scale bars, 10 cm in (a) and 1 cm in (b-g).

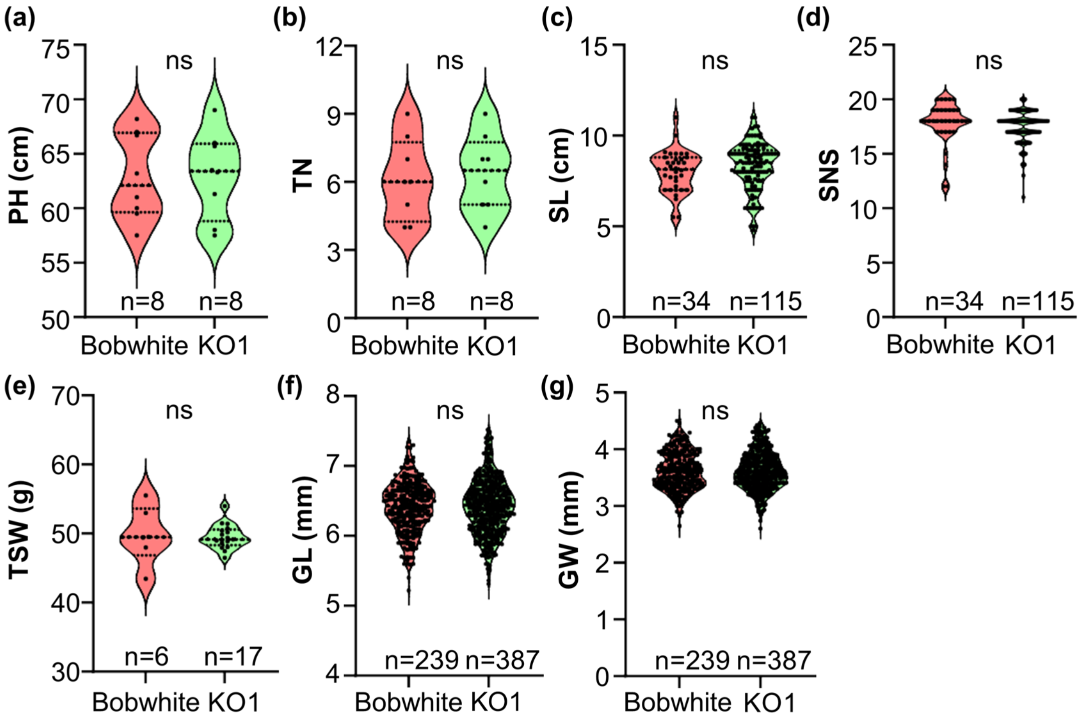


**Figure S9** Statistical analysis of agronomic traits in Bobwhite and *BGI1* KO1 mutant plants. (a) Plant height (PH); (b) Tillers number (TN); (c) Spike length (SL); (d) Spikelet number per spike (SNS); (e) Thousand-seed weight (TSW); (f) Grain length (GL); and (g) Grain width (GW). Plants were grown in growth chambers at 25℃ with a 16 h light / 8 h dark photoperiod. The violin plot shape reflects the distribution of each variable. Black circles represent single data points. The significance of differences was estimated using a two-sided unpaired *t*-test. ns, not significant ($P > 0.05$).

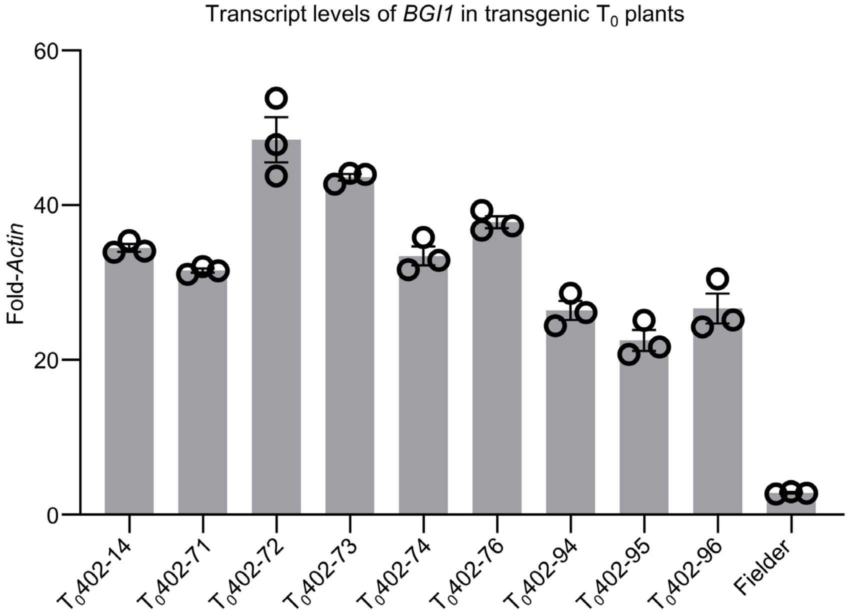


**Figure S10** Transcript levels of *BGI1* in overexpression transgenic $T_0$ plants. Transcript levels were measured using three technical replicates from a single plant and are expressed as fold-*Actin*. Error bars are standard errors of the means. Black open dots represent individual data points.

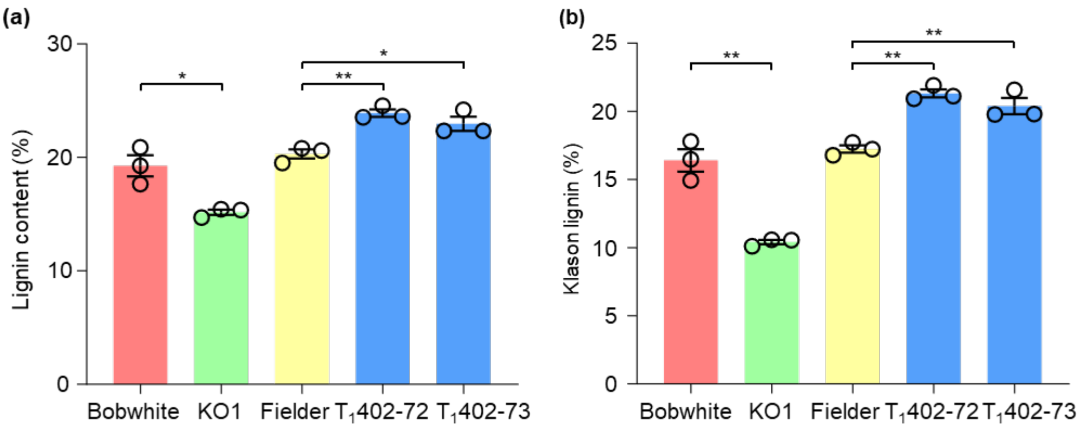


**Figure S11** Total lignin content and Klason lignin levels in stems. A comparison of total lignin content (a) and Klason lignin levels (b) in stems at the heading stage among WT, KO1 mutant plants, and OE transgenic lines ($T_1$402-72 and $T_1$402-73). Error bars are standard errors of the means (n = 3). *, *P* < 0.05; **, *P* < 0.01.

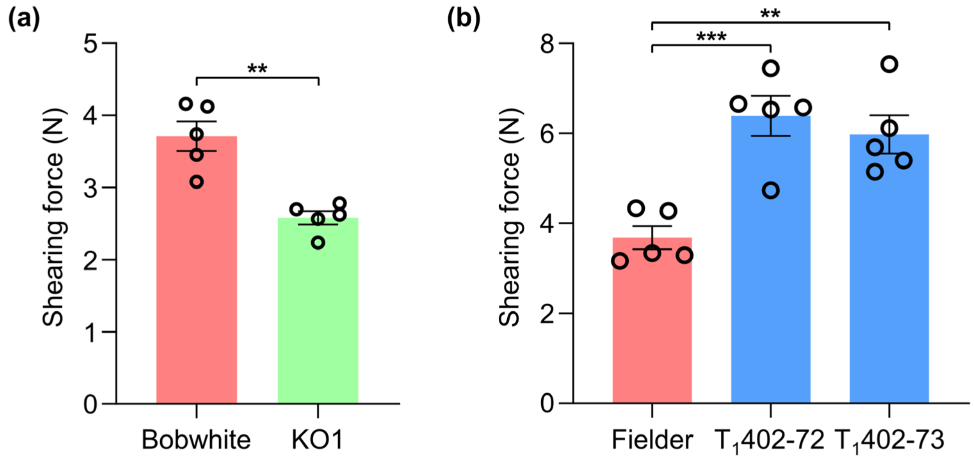


**Figure S12** Shearing force of WT, KO1 mutant plants, and OE transgenic lines ($T_1$402-72 and $T_1$402-73). (a) Shearing force of Bobwhite and KO1 mutant plants; (b) Shearing force of Fielder and OE transgenic lines. The stem shearing force was measured using a universal testing machine (Instron 5848 Microtester, Instron, USA). Data are presented as mean ± SEM. The significance of differences was estimated using a two-sided unpaired *t*-test. Black open dots represent individual data points. **$P < 0.01$; ***$P < 0.001$.

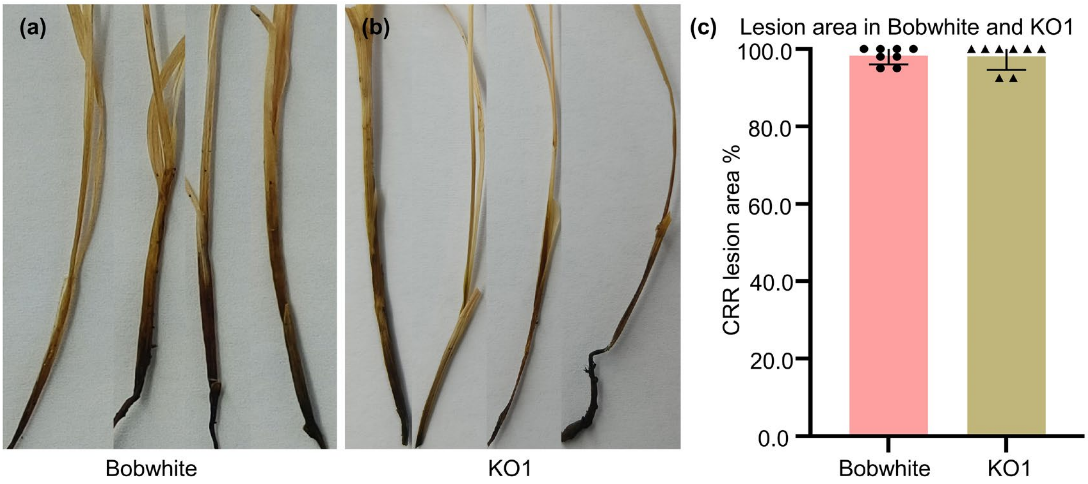


**Figure S13** Typical common root rot (CRR) symptoms and lesion areas in Bobwhite and KO1 mutant plants. (a-b) Phenotypic comparison of Bobwhite and KO1 mutant plants in response to CRR. (c) Lesion area comparison between Bobwhite and KO1 mutant plants in response to CRR. Error bars are standard errors of the means (n = 8). Lesion areas were determined from images of infected plants using ASSES v2 image analysis software. Black dots and triangles represent individual data points.

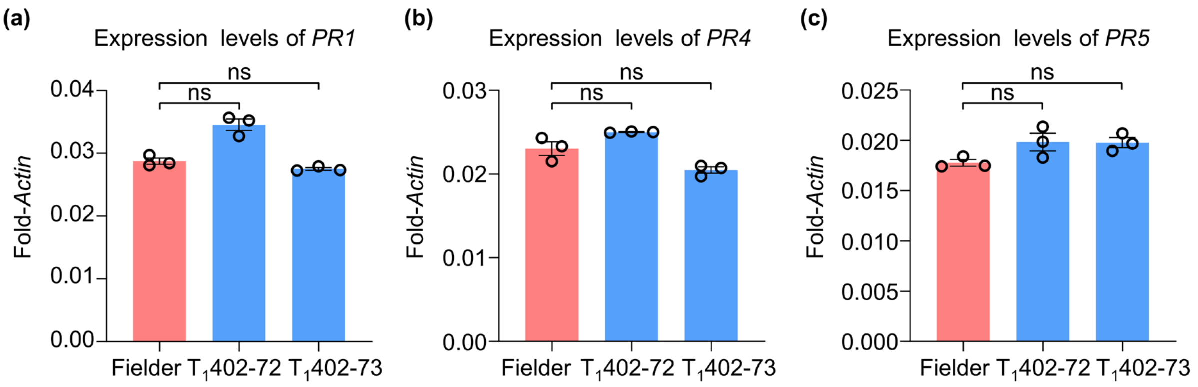


**Figure S14** Expression levels of pathogenesis-related (*PR*) genes in Fielder and *BGI1* overexpressing $T_1$ plants. (a) Expression levels of *PR1*. (b) Expression levels of *PR4*. (c) Expression levels of *PR5*. Stems at the heading stage from OE transgenic lines ($T_1$402-72 and $T_1$402-73) and Fielder were analyzed. Transcript levels were quantified relative to *Actin* using the $2^{-\Delta CT}$ method. Error bars indicate standard errors of the means (n = 3). Statistically significant differences were calculated using a two-sided unpaired *t*-test. Black open dots represent individual data points. ns, not significant ($P > 0.05$).

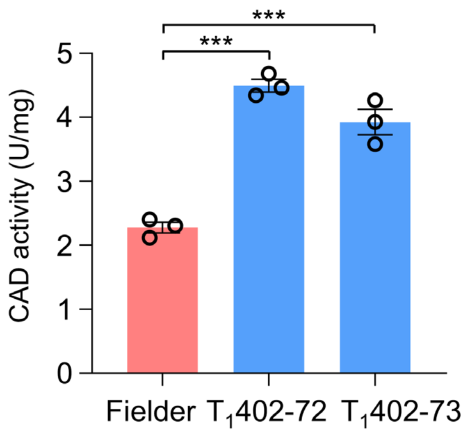


**Figure S15** Comparison of CAD activities between Fielder and OE transgenic lines ($T_1$402-72 and $T_1$402-73). Error bars are standard errors of the means (n = 3). Statistically significant differences were calculated using a two-sided unpaired *t*-test. Black open dots represent individual data points. ***, $P < 0.001$.

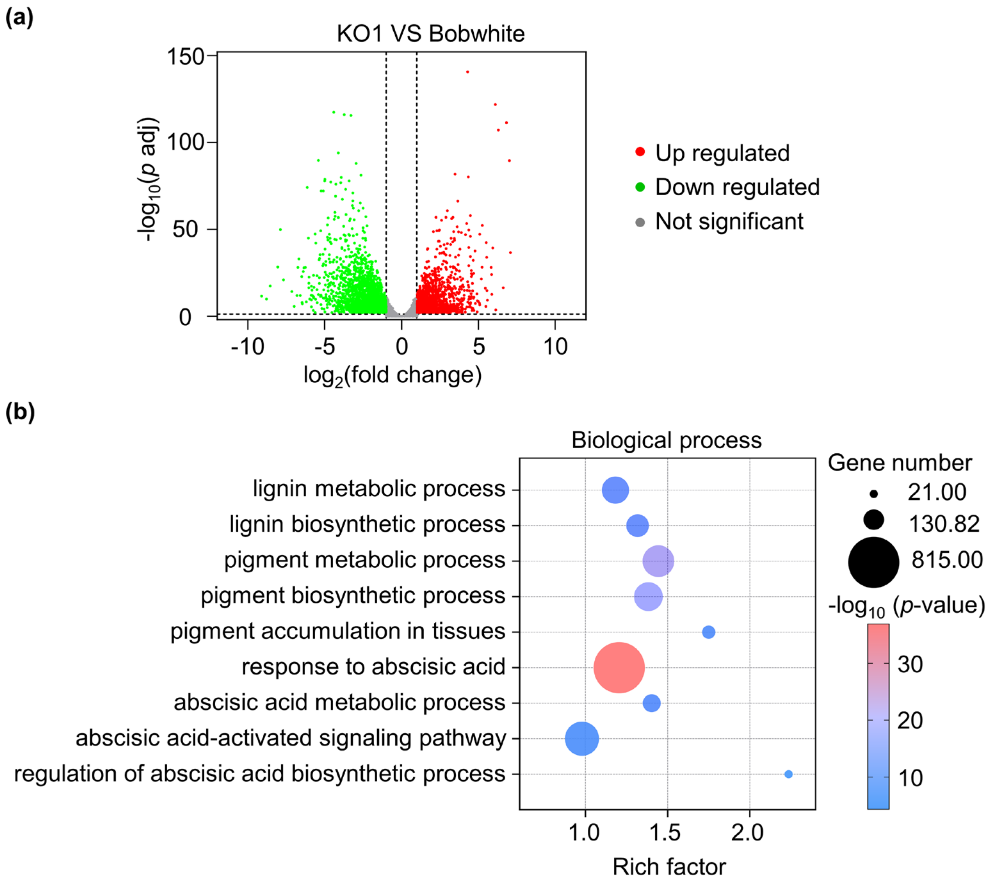


**Figure S16** Transcriptome analysis between KO1 mutant plants and Bobwhite. (a) Volcano plot displaying differentially expressed genes between KO1 and Bobwhite in the internodes at the heading stage. The x-axis represents the $\log_2$ (fold change) value, and the y-axis displays the -log10 (*p* adj) expression value. Red and green dots represent up-regulated and down-regulated genes (FDR < 0.01, *p*-value < 0.05, and $|\log_2$fold change$| > 1$) between KO1 and Bobwhite, respectively. (b) Major pathways of differentially expressed genes in biological process term enrichment analysis. The bubble size means the number of genes, while the bubble color corresponds to the -$\log_{10}$ (*p*-value). The adjusted *p* values in (a) were determined by the Benjamini-Hochberg multiple test correction. The unadjusted *p* values in (b) were determined using a two-sided Fisher's exact test, with no adjustments for multiple comparisons.

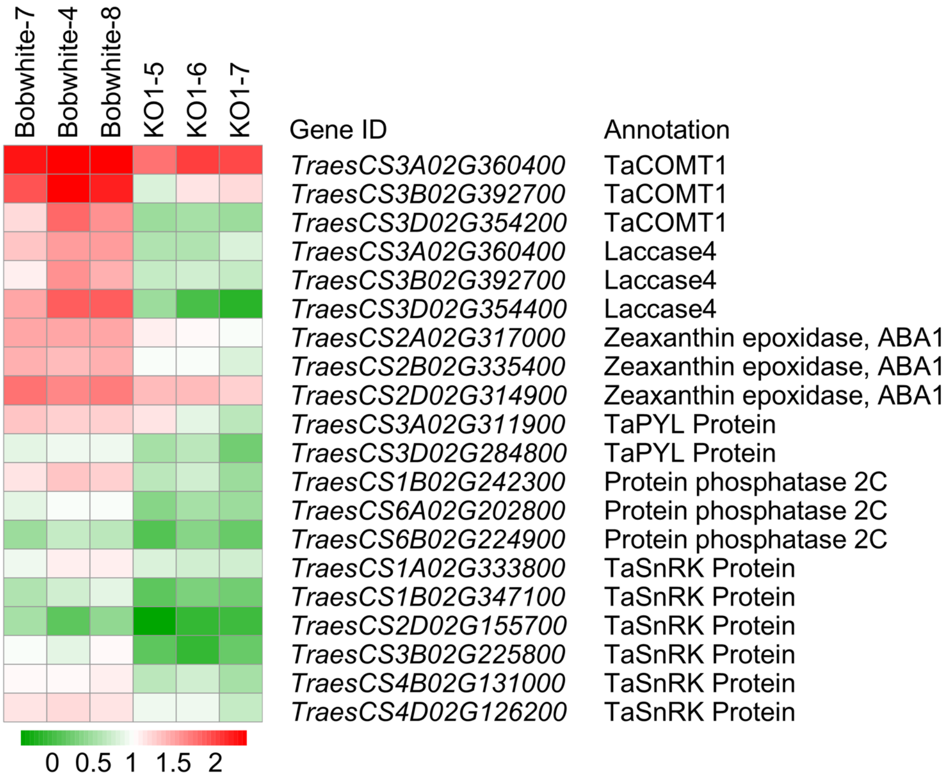


**Figure S17** Heatmap showing representative differentially expressed genes involved in lignin biosynthesis and ABA-related pathways. Differentially expressed genes between Bobwhite (Bobwhite-4, Bobwhite-7, and Bobwhite-8) and KO1 mutant plants (KO1-5, KO1-6, and KO1-7) were identified from RNA-seq data. Three independent biological replicates were analyzed for each genotype.

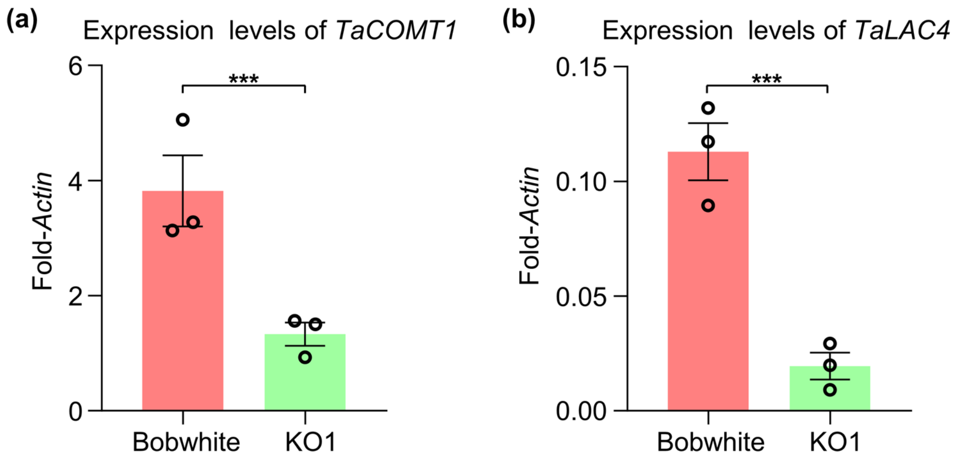


**Figure S18** Expression levels of lignin biosynthesis-related genes in Bobwhite and KO1 mutant plants. (a) Expression levels of *TaCOMT1* (*TraesCS3A02G534900*). (b) Expression levels of *TaLAC4* (*TraesCS3B02G392700*). Stems at the heading stage from KO1 mutant plants and Bobwhite were analyzed. Transcript levels were quantified relative to *Actin* using the $2^{-\Delta CT}$ method. Error bars indicate standard errors of the means (n = 3). Statistically significant differences were calculated using a two-sided unpaired *t*-test. Black open dots represent individual data points. ***$P < 0.001$.

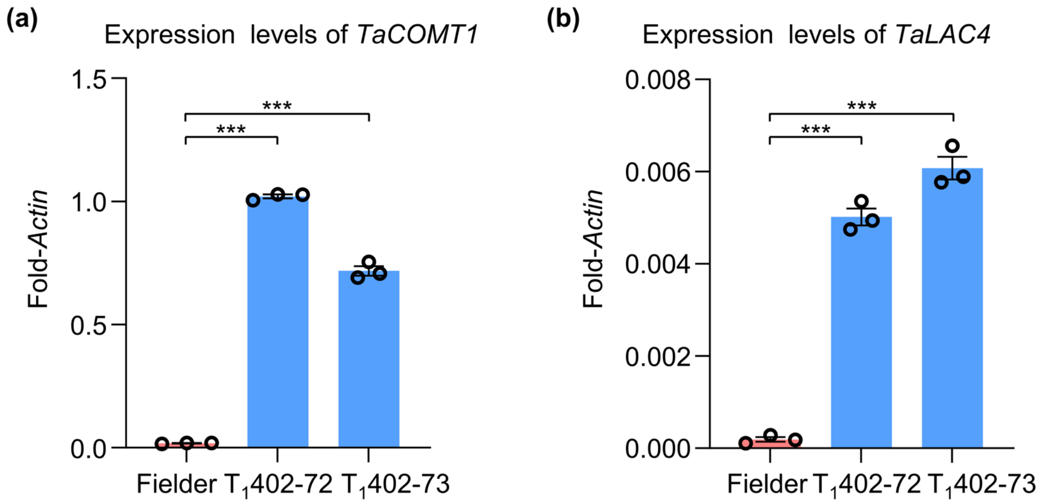


**Figure S19** Expression levels of lignin biosynthesis-related genes in Fielder and *BGI1* overexpressing $T_1$ plants. (a) Expression levels of *TaCOMT1* (*TraesCS3A02G534900*). (b) Expression levels of *TaLAC4* (*TraesCS3B02G392700*). Stems at the heading stage from OE transgenic lines ($T_1$402-72 and $T_1$402-73) and Fielder were analyzed. Transcript levels were quantified relative to *Actin* using the $2^{-\Delta CT}$ method. Error bars indicate standard errors of the means (n = 3). Statistically significant differences were calculated using a two-sided unpaired *t*-test. Black open dots represent individual data points. ***$P < 0.001$.

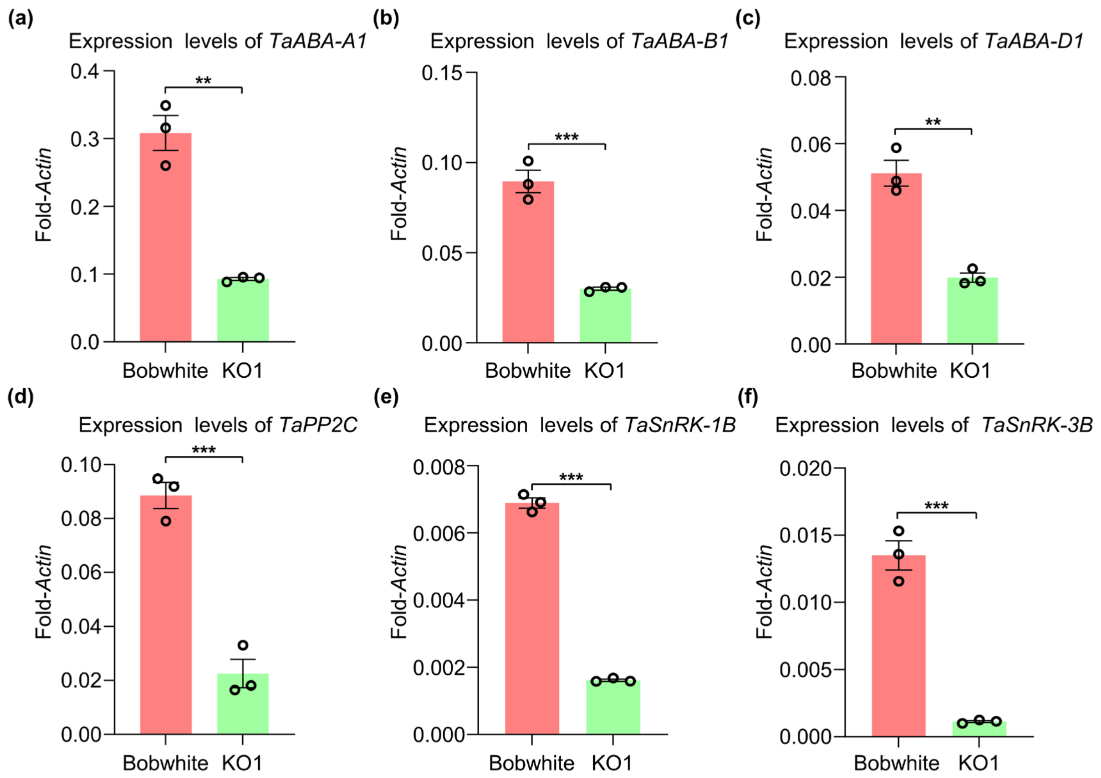


**Figure S20** Expression levels of ABA-related genes in KO1 mutant plants and Bobwhite. (a-c) Expression levels of *TaABA1* (*TraesCS2A02G317000*, *TraesCS2B02G335400*, and *TraesCS2D02G314900*). (d) Expression levels of *TaPP2C* (*TraesCS1B02G242300*). (e-f) Expression levels of *TaSnRK* (*TraesCS1B02G347100* and *TraesCS3B02G225800*). Transcript levels were quantified relative to *Actin* using the $2^{-\Delta CT}$ method. Statistically significant differences were calculated using a two-sided unpaired *t*-test. Error bars indicate means ± SEM (n = 3). Black open dots represent individual data points. ***$P < 0.001$.

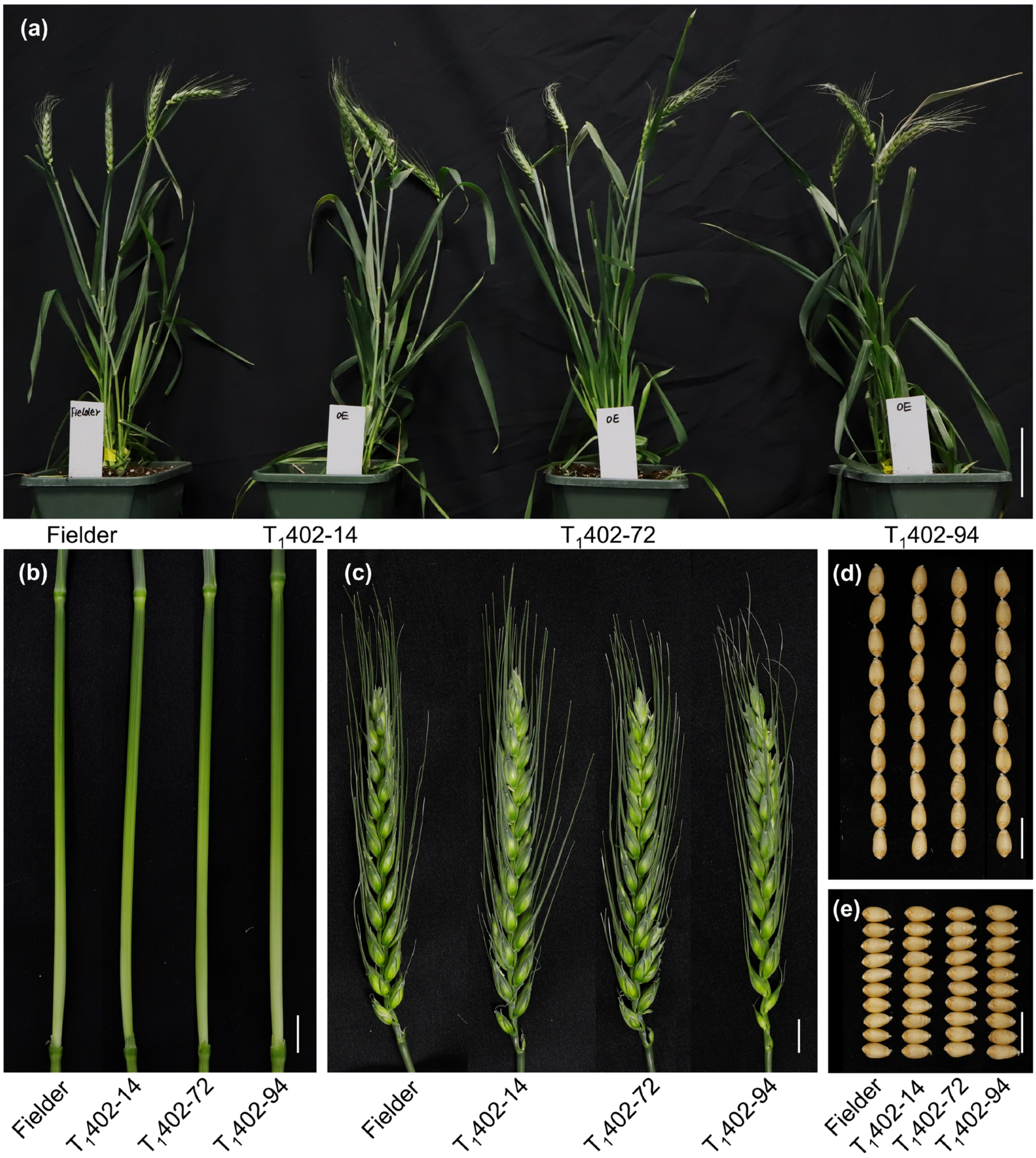


**Figure S21** Phenotypic comparison of Fielder and three *BGI1* overexpression transgenic lines. Close-up views of the plant (a), internode (b), spike (c), grain length (d), and grain width (e) between Fielder and three OE transgenic lines ($T_1$402-14, $T_1$402-72, and $T_1$402-94). Plants were grown in growth chambers at 25℃ with a 16 h light / 8 h dark photoperiod. Scale bars, 10 cm in (a) and 1 cm in (b-e).
